# Coinfection with chytrid genotypes drives divergent infection dynamics reflecting broad epidemiological patterns

**DOI:** 10.1101/2022.09.28.509987

**Authors:** Tamilie Carvalho, Daniel Medina, Luisa P. Ribeiro, David Rodriguez, Thomas S. Jenkinson, C. Guilherme Becker, Luís Felipe Toledo, Jessica Hite

## Abstract

By altering the abundance, diversity, and distribution of species — and their pathogens — globalization may inadvertently select for more virulent pathogens. In Brazil’s Atlantic Forest, a hotspot of amphibian biodiversity, the pet trade has facilitated the co-occurrence of previously isolated enzootic and panzootic lineages of the pathogenic amphibian-chytrid (‘Bd’) and generated new virulent recombinant genotypes (‘hybrid’). Epidemiological data indicate that amphibian declines are most severe in hybrid zones, suggesting that coinfections are causing more severe infections or selecting for higher virulence. We investigated how coinfections involving these genotypes shaped virulence and transmission. Overall, coinfection favored the more virulent and competitively superior panzootic genotype, despite dampening its virulence and transmission. However, for the least virulent and least competitive genotype, coinfection increased both pathogen virulence and transmission. Thus, by integrating experimental and epidemiological data, our results provide mechanistic insight into how globalization can select for, and propel, the emergence of introduced hypervirulent lineages, such as the globally distributed panzootic lineage of Bd.

## INTRODUCTION

Globalization can facilitate the emergence and spread of infectious pathogens (Daszak et al. 2001; Anderson et al. 2004). When hosts are exposed to completely novel pathogen species or genotypes, coinfections with multiple genotypes can allow new pathogen variants to emerge via recombination between genotypes (Brasier 2001; Stukenbrock et al. 2012). Genetically diverse or ‘mixed’ infections can also select for more virulent pathogens owing o within-host competition among different pathogen genotypes (Alizon et al. 2013).Competitively superior genotypes generally exploit host resources faster, and thus, limit the reproduction and transmission of prudent (less virulent) pathogens (de Roode et al. 2005). Moreover, even if more aggressive pathogens kill their hosts and reduce their own fitness, less virulent pathogens suffer a disproportionate disadvantage (linked to reduced transmission) and may be eliminated by natural selection (May & Nowak 1995, Read & Taylor 2001, Alizon et al. 2013). However, empirical studies examining genetically diverse infections in wild, non-laboratory host-pathogen systems remain limited. A central research objective, therefore, is to understand how mixed-genotypes infections affect disease outbreaks in the wild, which will enable the design of more effective public health and conservation interventions (Brasier 2001; Stukenbrock et al. 2012).

In Brazil’s Atlantic Forest, a hotspot of amphibian biodiversity (Toledo et al. 2021), globalization has led to the co-occurrence of enzootic (Bd-Asia-2/Brazil) and panzootic (Bd-GPL) lineages of the amphibian-killing fungus *Batrachochytrium dendrobatidis* (Bd), which have different introduction histories. In addition to these two strains, novel recombinant genotypes (hereafter ‘hybrids’) have also emerged (Jenkinson et al. 2016; O’Hanlon et al. 2018; Fisher & Garner 2020). Bd is a cutaneous fungus that disrupts key physiological functions of amphibian skin (Voyles et al. 2011), leading to the potentially fatal disease chytridiomycosis (Scheele et al. 2019). Bd genotypes from the enzootic and panzootic lineages vary in phenotypic traits, competitive ability, and virulence (Lambertini et al. 2016; Becker et al. 2017; Jenkinson et al. 2018; Greenspan et al. 2018). In addition, hybrid genotypes, such as the ones resulting from the recombination between Bd-Asia-2/Brazil and Bd-GPL, can be more virulent relative to their parental lineages; this was observed in one host species during an experimental trial using single-genotype infections (Greenspan et al. 2018). The co-occurrence of enzootic, panzootic and hybrid genotypes of Bd in Brazil’s Atlantic Forest suggests that coinfections among these genotypes carry important implications for disease dynamics and virulence evolution (O’Hanlon et al. 2018; Byrne et al. 2019).

Notably, in Brazil, most amphibian declines have occurred in hybrid zones; locations where Bd-Asia-2/Brazil, Bd-GPL, and hybrid genotypes co-occur (Jenkinson et al. 2016; Carvalho et al. 2017). This observation suggests that coinfections may either lead to more severe infections or select for more virulent lineages. Therefore, it is crucial to understand whether coinfections could shape competitive interactions among these genotypes and alter transmission and virulence (*e*.*g*., Jenkinson et al. 2018).

Here, we examined how coinfection between four divergent genotypes of Bd from two distinct lineages and a hybrid affect competitive interactions within-hosts. Then, we asked whether these interactions altered pathogen virulence (host survival and lifespan) and transmission potential (using Bd load as a proxy for future transmission potential). Overall, coinfection favored the most virulent genotype (P1), despite decreasing its transmission potential and virulence. However, coinfection increased the virulence of the least competitive and least virulent genotype (P2). These experimental results, combined with our field data, provide mechanistic insight into how the within-host competitive interactions could lead to the distribution pattern of Bd lineages across Brazil’s Atlantic Forest.

## METHODS

We used five Bd genotypes from strains that vary in their degree of relatedness, competitive ability (using early establishment as a metric), virulence, and transmission potential (total Bd load) to examine how coinfection affects disease outcomes. We performed analyses for each genotype alone or when in mixed-infections with one or two other genotypes. We used genotypes from the most recently derived lineage of Bd, the Global Panzootic Lineage (Bd-GPL; Fig. 1A; hereafter, ‘P1’ or ‘P2’), as the reference genotypes that we placed in competition with two enzootic (‘E1’ and ‘E2’) and a hybrid genotype (‘H’). We use P1 and P2 as reference in an effort to capture the arrival of Bd-GPL in Brazil, where coinfections with the local enzootic lineage led to the emergence of the hybrid genotype.

**Figure 1.**
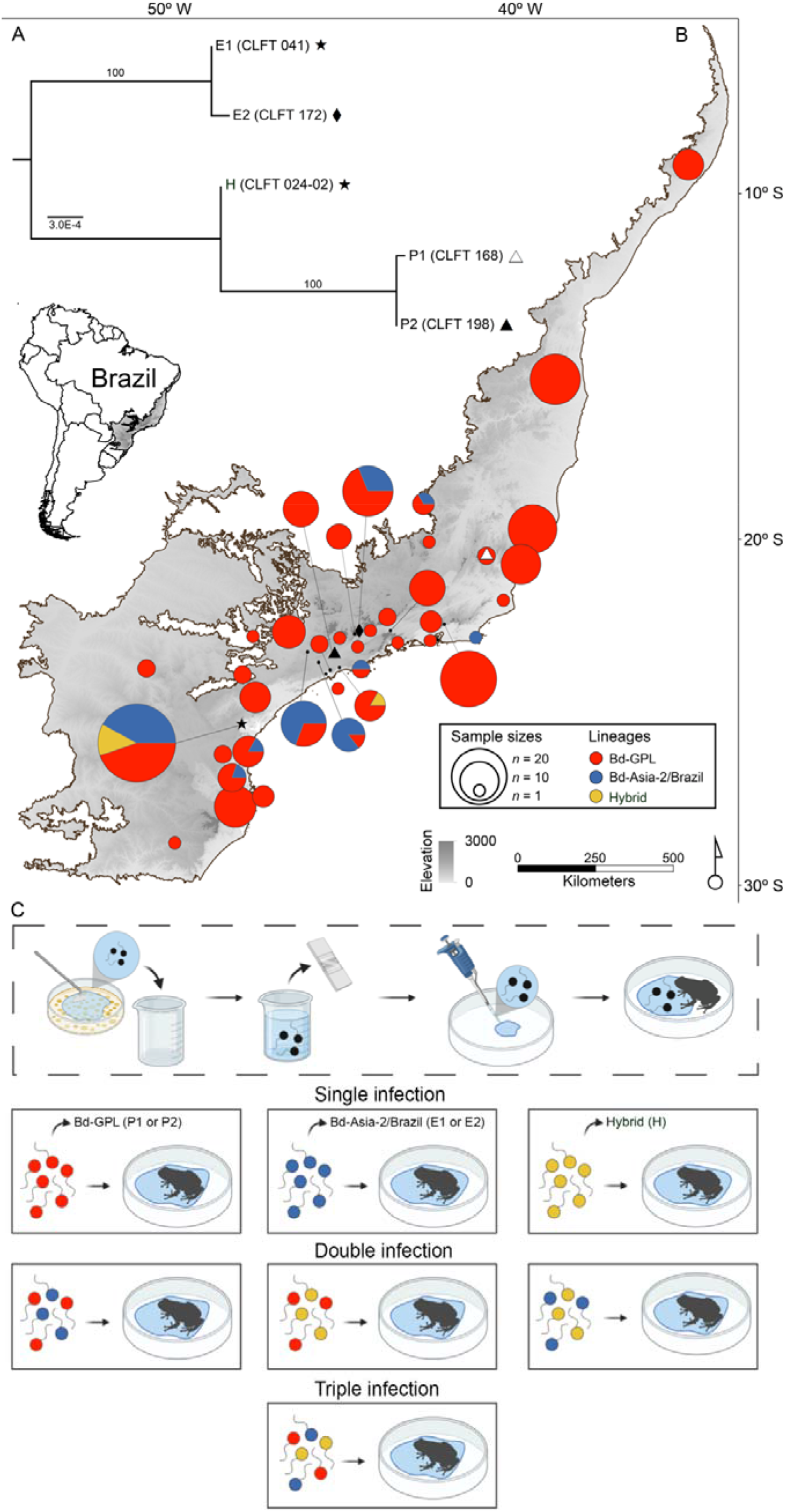
Phylogenetic relationships among *Batrachochytrium dendrobatidis* (Bd) genotypes, geographical distribution of their lineages, and experimental design for single- and mixed-genotype infections. (*A*) Phylogenetic tree of Bd genotypes used in the challenge assay. (*B*) Spatial distribution of Bd lineages or hybrid genotypes identified from multiple amphibian species across the Brazilian Atlantic Forest (highlighted area in map) (Table S5). On map, symbols represent the location where we isolated the pathogen genotypes used in this experiment. White triangle: P1, black triangle: P2, diamond: E2, star: E1 and H. (*C*) Infection assay procedure. In order to prepare the Bd solutions, we flooded Bd cultures with distilled water, quantified Bd zoospore density using a hemocytometer, and adjusted all solutions to the same concentration (see Methods for additional experimental details).

To examine whether observed within-host competitive outcomes in the experiment reflect distribution patterns of Bd lineages across Brazil’s Atlantic Forest, we genotyped Bd isolates obtained from tadpoles of natives and introduced (i.e., *Aquarana catesbeiana*) frog species sampled at multiple locations (Fig. 1B, Table S1). The sampling of tadpoles from native species took place in natural ponds and streams, and the sampling of *A. catesbeiana* tadpoles took place in farms where they are bred for commercial purposes. We determined tadpoles infection status by examining the degree of depigmentation in the oral disc (Knapp & Morgan 2006). In addition, we complemented our field data by including previously published results of Bd genotyping from adults and tadpoles sampled in the study region (Table S1). In total, this dataset included 237 isolates (182 tadpoles and 55 adults) collected from 42 species between 1987 and 2016.

The isolation, culturing, DNA extraction and genotyping procedures used in all Bd isolates included in this study (experiment and field collection) follow those described in Jenkinson et al. (2016).

### Infection assay

#### Pathogen genotypes and amphibian host

The Bd isolates used in our infection assays were obtained from tadpoles of captive North-American bullfrogs, *A. catesbeiana* (P2 and E2) and wild anurans (P1, E1 and H) (Table S2).

Prior to the experiment, isolates were maintained at 4 ºC, passaged every four months, and their number of passages range from 4 to 20 (Table S2). Although virulence attenuation can occur in Bd isolates serially passaged for long periods (Langhammer et al. 2013; Refsnider et al. 2015), Bd isolates with as much as 50 passages (compared to isolates with 10 passages) can present higher rates of zoospore production and overall population growth, and induced higher infection loads and prevalence in inoculation experiments (Voyles et al. 2014). Thus, we infer that the number of passages did not, at least to a noticeable level, affect probability of Bd colonization and reproduction in our assay.

To verify the phylogenetic relationships between our isolates in the challenge assays, we inferred a tree using published sequence data for these Bd genotypes (*SI Appendix*). Using six MLST marker sequences (8009×2, BdC24, BdSC4.16, R6046, BsSC6.15, BdSC8.10) previously generated from associated studies (Jenkinson et al. 2016; Ribeiro et al. 2019), we constructed a hetequal distance dendrogram (James et al. 2009; Jenkinson et al. 2016) using PAUP* version 4.0b10 (Fig. 1A). The support values for the major clades (Bd-GPL and Bd-Asia-2/Brazil) were estimated by bootstrapping over 100 replicates.

We used the terrestrial frog *Eleutherodactylus johnstonei* (Anura; Eleutherodactylidae), which is a direct-developing species (*i*.*e*., lacking a larval stage and with embryo hatching as froglets), native to the Lesser Antilles (Schwartz 1967), that was introduced to the city of São Paulo, Brazil, at least 27 years ago (Toledo & Measey 2018). Frogs used in the experiment were collected from this introduced population.

*Eleutherodactylus johnstonei* is known to tolerate Bd infections and is considered a potential reservoir species in Dominica and Montserrat (Hudson et al. 2019). However, in Brazil, the introduced population of *E. johnstonei* have remained restricted to a small urban habitat and free of Bd. Two recent studies, after testing 100 frogs in total, did not detect Bd at this location (Forti et al. 2017; Mesquita et al. 2017). Besides, this host population exhibited high levels of susceptibility to Bd in a previous experiment (Mesquita et al. 2017), therefore serving as a good model-species for our experiments.

At the collection site, we selected adults with approximately the same size (SVL mean = 21.12 ± sd 2.66 mm, n = 95), and handled each adult frog with a new pair of disposable gloves to avoid potential Bd cross-contamination, and gently placed them into individual containers. We then transported the frogs to the laboratory in refrigerated coolers. To confirm that all hosts were Bd-free prior to experimental inoculations, we swabbed the skin of all the frogs upon arrival at the laboratory following a standard protocol (Hyatt et al. 2007). We stored swabs at −20 ºC until processing.

After swabbing, we housed the frogs individually in plastic enclosures (22 × 15 × 8 cm), with a layer of sphagnum moss that was previously autoclaved and moistened with distilled water. Frogs were housed in temperature-controlled rooms at 20 ºC on a 12-hour day-night cycle. We monitored frogs twice a day throughout the experiment and fed them calcium-enriched crickets *ad-libitum* twice a week.

#### Experimental exposure

Experimental treatments included single-genotype exposures and mixed genotype exposures with either two or three genotypes with all possible inter-lineage combinations (Fig. 1C). In total, the experiment included 17 Bd exposed treatments and one control group for a total of 297 hosts of the invasive species *E. johnstonei* (Table S3).

To prepare Bd suspensions for the challenge assay, we transferred liquid cultures of each genotype to individual agar plates containing 1 % tryptone and allowed them to grow for eight days at 11 ºC under dark conditions. We then collected zoospores of each Bd genotype by flooding culture plates with 2 ml of distilled water for 45 min to induce zoospore release and then gently scraping the surface to maximize collection. From each zoospore suspension, we prepared standard inoculation solutions (concentration 2.98 × 10^6^ zoospores / ml) by collecting a 1 ml subsample and quantifying the zoospore density using a hemocytometer (Fig. 1C).

For pathogen exposure, individual hosts were placed in Petri dishes with 1.5 ml of zoospore suspension containing a fixed inoculum dose (2.98 × 10^6^ zoospores / ml), and were exposed for 45 min at 20 ºC (Fig. 1C). Frogs from the control group were exposed to 1.5 ml of distilled water. Individuals from coinfection treatments (those exposed to two or three genotypes) were exposed to a zoospore suspension containing equal proportions of each Bd genotype (final volume of 1.5 ml). To quantify Bd infection following exposure, we collected skin swabs on day 21 post exposure and after death or at the end of the experiment (day 76) following a standard protocol (Hyatt et al. 2007). Both collection days span multiple Bd life cycles (Berger et al. 2005). Swabs were extracted using the DNeasy blood and tissue kit (Qiagen, Valencia, CA, USA) following the manufacturer’s protocol, with a minor modification that consisted of an extended incubation time (four hours) in the lysis step to increase DNA yield.

#### Genotyping of coinfections

To simultaneously detect Bd lineages (and hybrid) and estimate infection loads during experimental coinfections, we developed a genotyping assay for this study. Overall, we leveraged two genotyping assays to detect the presence or absence of Bd-Asia-2/Brazil, Bd-GPL, and the hybrid (Schloegel et al. 2012) in a mixed sample. Assay Bdmt_26360 targets a mitochondrial SNP and distinguishes between Bd-Asia-2/Brazil (allele A, 6-FAM) and Bd-GPL or hybrid (allele G, VIC) (Jenkinson et al. 2018). When allele G was detected in a mixed sample by the mtDNA assay, then we performed a second genotyping reaction using assay BdSC9_621917_AC (present study; Table S4) to target a SNP in the nuclear genome to determine whether the hybrid genotype (allele C, 6-FAM) was present or absent. Absence of allele C would indicate only Bd-GPL was present in the sample (Fig. S1). This nuclear assay allowed us to accurately distinguish between Bd-GPL and hybrid in 100% of the frogs from the double infection and over 80% from the triple infection treatments.

We used the skin swab DNA extractions as input for qPCRs in 20 μl volumes composed of 6 μl of the sample template, 10 μl of TaqMan Fast Advanced 2X Master Mix (Applied Biosystems, Inc.), 3 μl of nuclease-free water, and 1 μl of the SNP assay mix (Applied Biosystems, Inc.), at 20X concentration (18 μM forward primer, 18 μM reverse primer, 4 μM 6-FAM probe, and 4 μM VIC probe). On a QuantStudio 3 (Applied Biosystems, Inc.), cycling conditions consisted of 95 °C for 20 s, then 95 °C for 1 s and 60 °C for 20 s (data collection step) for 50 cycles. We analyzed amplification curves for either VIC or 6-FAM using the Standard Curve application that is part of the Thermo Fisher Connect cloud-based software.

#### Bd *quantification using standard curves*

We sourced three strains from the Collection of Zoosporic Eufungi at the University of Michigan to serve as genotyping positive controls and quantification standards. These included previously characterized isolates CLFT 041 (Bd-Asia-2/Brazil), CLFT 095 (Bd-GPL), and CLFT 024-02 (hybrid), of which two (CLFT 041 and CLFT 024-02) were used in the infection assay. Each isolate was transferred to 10 ml of 1 % Tryptone liquid media and grown at 17 °C for 14 days in cell culture flasks. After confirming the presence of active zoospores, the medium was agitated and 1 ml was transferred to 1.5 ml centrifuge tubes that we spun at 10,000 rcf for 10 min to pellet the zoospores. The liquid was decanted off and the pellets were used as input for the GenJet DNA extraction kit (Thermofisher Inc.) following the manufacturer’s recommendations but using 100 μl final elution volumes. Replicate tubes were pooled and the resulting purified DNA was quantified on a Qubit 3 (Applied Biosystems). DNA for each isolate was normalized to 1.34 ng/μl. Serial dilutions with a dynamic range of 1.34 × 10^−1^ to 1.34 × 10^−5^ ng/μl were used as inputs for qPCR standards for each strain type and assay. Then, genotyping amplification curves were used to estimate Bd load measured in ng/μl based on the strain specific standard slopes (Table S5; see Longo et al. 2013). For statistical analyses, we used Bd measures (represented by quantified DNA concentrations) from the mitochondrial assay (Bdmt_26360), and transformed them to pg/μl to facilitate the calculation of parameter estimates and interpretation of analyses.

### Statistical analyses

To ensure that our infection methodology was consistent across treatments, we first quantified the proportion of hosts that become infected over the course of the experiment using Generalized Linear Models (GLMs) with binomial errors (the proportion of hosts that became infected did not differ across genotypes or coinfections; Fig. S2 A-C; Binomial GLM, all *p*-values > 0.05). Then, we tested for differences in competitive ability (using early establishment of Bd as a metric – 21 days post-inoculation), virulence (lifespan and survival), and transmission potential (using total Bd load – after death or on day 76) of each genotype when alone and when in competition with either one or two other genotypes using a combination of generalized linear models (GLMs), planned *a priori* orthogonal contrasts, and survival analyses.

For GLMs, we chose the most appropriate distribution for each model using the R package *fitdistrplus*. For virulence and transmission potential, we included only the animals hat were infected (and excluded animals that were exposed but uninfected; n = 3: E1+H = 1, P2+H = 1, P2+E2+H = 1). To understand how coinfections affected key traits of the reference genotypes, we used planned *a priori* orthogonal contrasts (Gotelli & Ellison 2004, Bolker 2008, Crawley 2012).

We computed survival curves using the Kaplan-Meier method, which was implemented using the *survfit* function from the package *survival* (Therneau 2021), and compared survival curves by conducting log rank tests using the function *survdiff* (also from the package survival). We conducted *post-hoc* comparisons of survival curves using the function *pairwise_survdiff* from the package *survminer* (Kassambara et al. 2019), which computes adjusted *p*-values to correct for false discovery rate (we used the method ‘BH’, Benjamini & Hochberg 1995). We examined the effect of coinfections by comparing survival curves of Bd genotypes from the panzootic lineage in single-genotype infections (P1 or P2) with their respective mixed infection treatments, which included the Bd genotypes from the enzootic lineage (E1 and E2) and the hybrid (H). All statistical analyses were conducted in R statistical software version 1.4.11 (R Core Team 2020).

## RESULTS

We found links between genotype virulence and competitive ability, but the direction of this relationship varied across particular Bd genotypes and their introduction history (*i*.*e*., enzootic *vs*. panzootic). The most competitive genotype, the panzootic P1, was also the most virulent genotype whereas the enzootic and hybrid genotypes were less competitive than the panzootic genotypes and showed intermediate virulence.

The panzootic Bd genotype P1 was the fastest to establish and outgrow all other genotypes in single-genotype infections (Bd load on day 21 of P1 to all other genotypes: *p* < 0.001; Fig. 2A). However, P1 suffered competitive suppression. In all mixed infections nvolving P1, early establishment was suppressed relative to the single infection trials (GLM single/mixed: *p* < 0.001; Fig. 2B). For P1, the degree of competitive suppression – at early stages of infection – was strong in triple infections involving E1 but relatively weak in triple infections with E2 (Fig. 2A, B). These results underscore that virulence evolution may depend strongly on genotype-specific traits.

**Figure 2.**
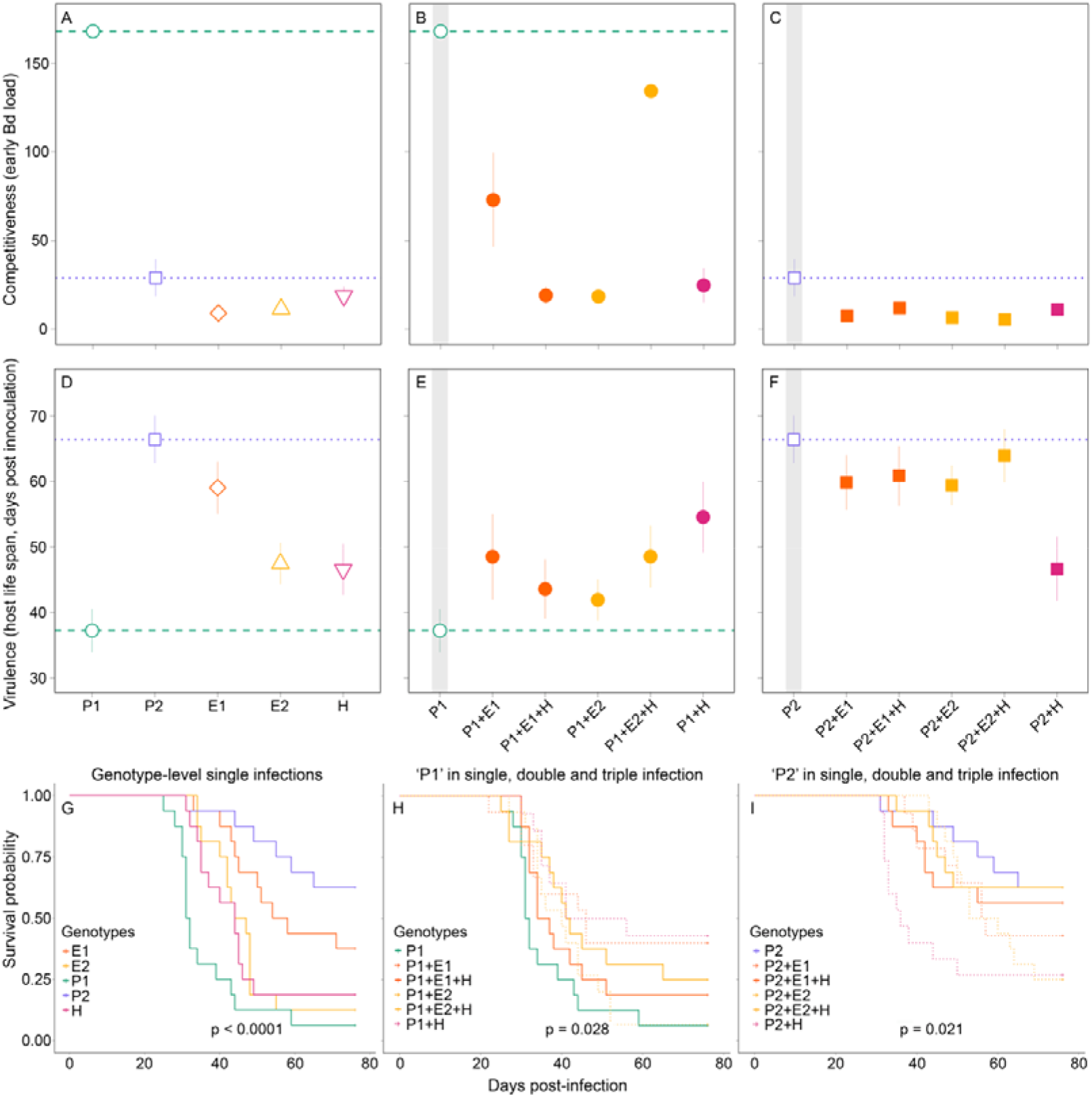
Competitive interactions and virulence vary across hosts infected with single- and mixed-genotypes of *Batrachochytrium dendrobatidis* (Bd). Grey bars highlight single-genotype infections. (*A-C)* Competitiveness of each genotype is represented as the average Bd load (in pg/μl) during the first time point of the assay (day 21 postinoculation). (*D-F*) Virulence measured as host-lifespan and (*G-I*) survival probability. The presence of multiple genotypes reduced the virulence (*i*.*e*., increased host life-span and survival) of the most virulent genotype (P1). Coinfections with the least virulent genotype (P2), however, did not influence host life-span, though they had an effect on survival or median survival time. For survival curves, *p*-values are from Log-rank tests.

Competitive suppression also reduced the virulence and transmission potential of P1. In single-genotype infections, P1 was more virulent than in mixed-genotype infections in both mean lifespan (GLM single/mixed: *p* = 0.052; Fig. 2D *vs*. 2E) and survival (Log-rank test: X^2^ = 12.50, df = 5, *p* = 0.03; Fig. 2G-H). In most cases, host survival rates slightly increased when coinfected. This trend was evident in the two-genotype infection with the hybrid genotype, although this trend was not statistically significant (median survival time P1: 31.5 days; P1+H: 48.5 days; adjusted-*p* = 0.09).

These longer lifespans, however, did not always lead to an overall increase in transmission potential. In all coinfections involving P1, transmission potential (total Bd load) was lower than in single-genotype infections (GLM single/mixed: *p* = 0.025; Fig. 3A, B). Thus, even though coinfected hosts lived longer, coinfections with the enzootic genotypes and hybrid still suppressed transmission potential for the most virulent genotype (P1) relative to the single-genotype infection.

**Figure 3.**
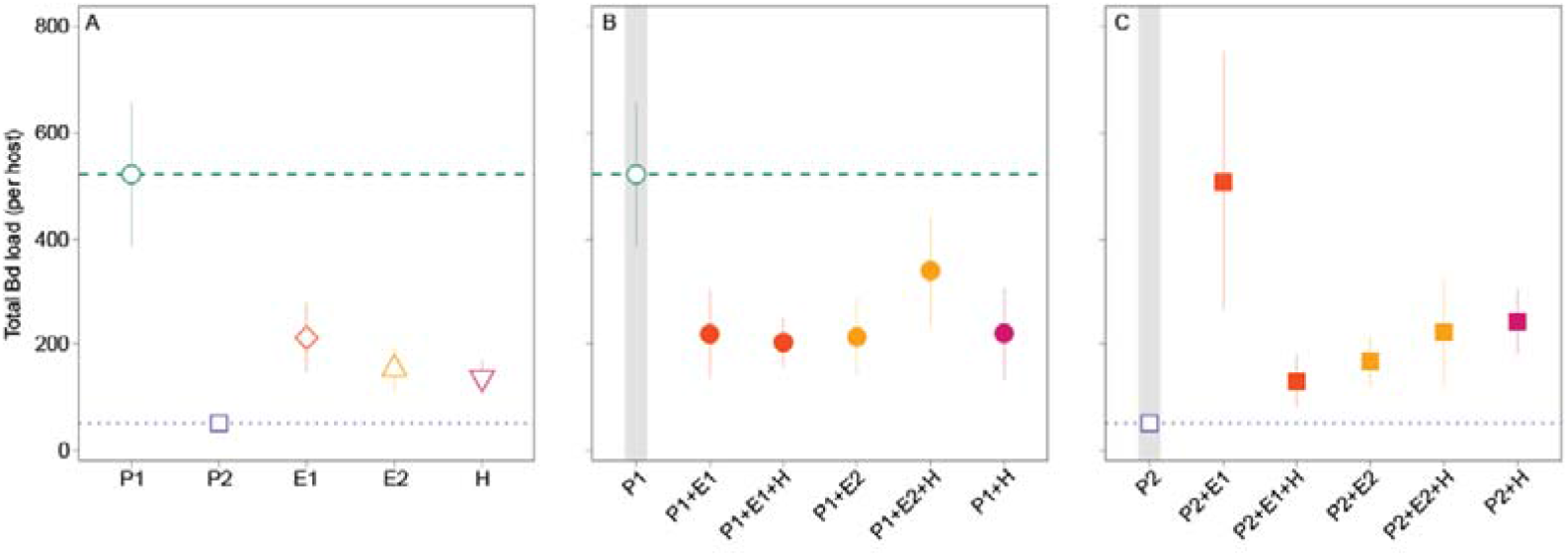
Transmission potential of *Batrachochytrium dendrobatidis* (Bd) in both single- and mixed-genotype infections. (*A-C)* Transmission potential is represented as the average Bd load (in pg/μl) after death or on day 76 at the end of the experiment. Grey bars highlight single-genotype infections. (*A* and *B*) Competitive suppression could either help reduce the transmission potential of more virulent genotypes (P1) or (*A* and *C*) increase the transmission potential of less virulent genotypes (P2).

In contrast, P2 was slower relative to P1 to establish and replicate in single infections and therefore, had a low competitive ability (Fig. 2A). In mixed infections with P2, it is likely that facilitation occurred for two reasons. First, the total Bd load (transmission potential) was higher in mixed infections relative to single-genotype infections (GLM single/mixed: *p* < 0.001; Fig. 3). For E1 in particular, coinfections with P2 lead to drastic increases in total Bd loads with levels reaching those similar to P1 single-genotype infections (Fig. 3A-B *vs*. C). Second, the proportion of P2 in these mixed infections was substantially lower relative to either the E1 or E2 genotypes (below 0.25, Fig. 4A). In triple infections, however, this facilitation was hindered by the hybrid genotype (H), which competitively suppressed the more virulent genotype but enhanced the less virulent genotype (Fig. 3A *vs*. C and Fig. 4A *vs*. C).

**Figure 4.**
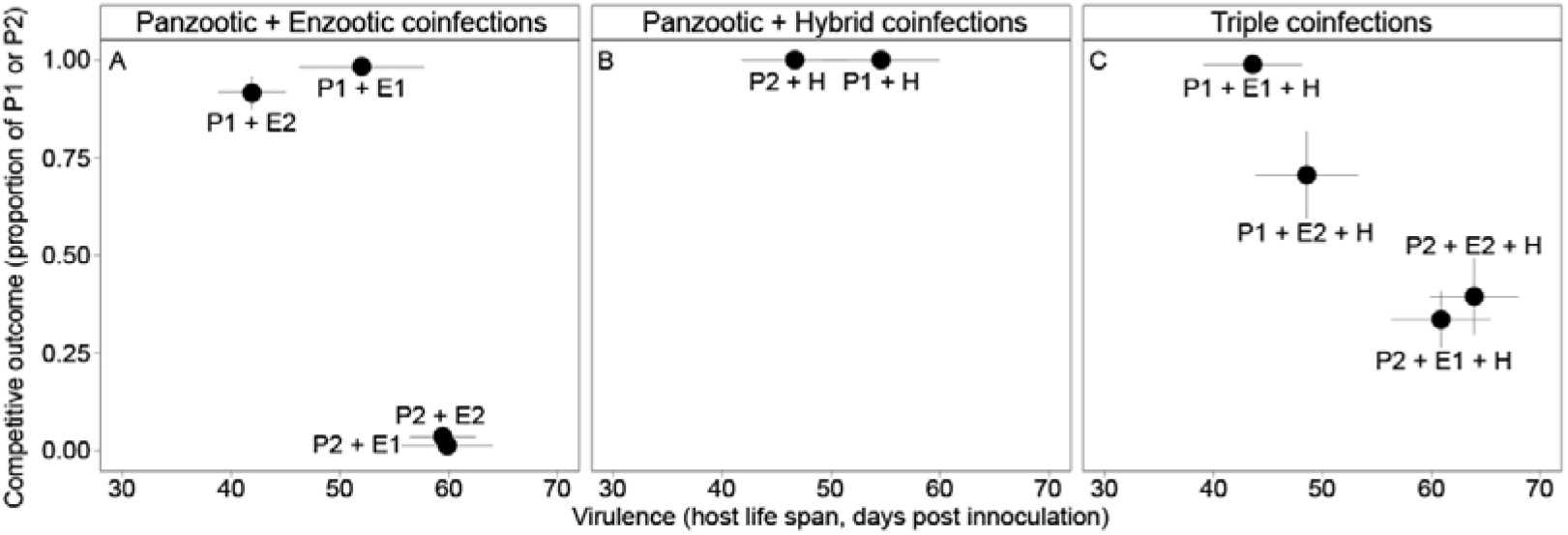
Competitive outcomes in mixed-genotype infections of *Batrachochytrium dendrobatidis* (Bd). (*A-C*) Competitive outcomes for the mixed-genotype infections were based on the proportion of the total pathogen load (in pg/μl) that was either the most and least virulent genotype (P1 and P2, respectively).

Importantly, despite the increase in pathogen loads (Fig. 3), coinfections involving P2 did not show substantially increased virulence (Fig. 2). For instance, mixed infections with P2 had no effect on mean lifespan (GLM single/mixed: *p* = 0.113; Fig. 2F) but slightly decreased host survival rates (Log-rank test: X^2^ = 13.30, df = 5, *p* = 0.02; Fig. 2G, I).

As for the Bd isolates collected from native species in the wild and *A. catesbeiana* individuals from frog farms, we determined that 80% (n = 190) of the isolates belong to the Bd-GPL lineage, followed by Bd-Asia-2/Brazil with 17% (n = 41) and hybrid genotype with 2% (n = 5) (Fig. 1B).

## DISCUSSION

Coinfections by multiple pathogen genotypes are common in humans (reviewed in Balmer & Tanner 2011), plants (Hood 2003; Susi et al. 2015), and wildlife (Rauch et al. 2005; Pickering et al. 2000). Theory predicts that coinfections have the potential to fundamentally alter infectious processes within- and between-hosts and thus carry important implications for epidemiological dynamics and virulence evolution (reviewed in Alizon et al. 2013). We show that coinfection can have differential outcomes on pathogen load and transmission potential over the course of infection depending on the combination of pathogen genotypes and their introduction histories. For a highly competitive and virulent genotype, such as P1 from the Bd-GPL lineage, these within-host interactions among pathogen genotypes reflect both global (O’Hanlon et al. 2018) and regional patterns of Bd outbreaks.

We found that some genotypes are competitively suppressed during coinfections, while other genotypes benefit from coinfections. For the most virulent and competitive genotype (P1), competitive suppression during coinfection reduced both virulent effects on host survival and transmission potential (total Bd load). The decline in transmission potential was likely driven through competitive suppression such that even though coinfected hosts lived longer, which could lead to higher shedding rates, overall pathogen production was reduced relative to hosts infected with a single pathogen genotype. For less virulent genotypes, however, facilitative interactions among pathogen genotypes had little effect on virulence but did increase transmission potential. Thus, while highly virulent genotypes may be subject to competitive suppression during coinfections, less competitive and virulent genotypes may facilitate other genotypes leading to increased transmission potential.

Links between competitiveness and virulence in the genotype P1 are consistent with global epidemiological patterns of this pathogen. Specifically, Bd-GPL is both globally distributed and associated with mass mortalities of amphibian hosts (James et al. 2015; Lips et al. 2016). Additionally, our results align with a previous study that demonstrated that Bd-GPL was more competitive than the enzootic lineage in the first four weeks post-inoculation (Jenkinson et al. 2018), a critical phase for successful pathogen colonization.

However, unlike P1, which was isolated from a wild host, the less competitive and virulent genotype P2 was isolated from a bullfrog farm. Such controlled environments with high density and low diversity of hosts can favor the emergence of highly virulent genotypes by increasing transmission potential (Frank 1996). However, the lower competitive ability, virulence and transmission potential observed in P2 may, in contrast, indicate that conditions associated with farms can also select for less virulent genotypes, possibly due to a constant availability of hosts resources. In addition, a high and constant availability of hosts may also allow pathogens to reallocate resources to suppress the host immune system. Thus, the evolution of pathogen virulence under farm conditions may be complex and difficult to predict, as suggested by Ribeiro et al. (2019) in a study where distinct Bd genotypes with different degrees of virulence were isolated from bullfrog farms.

Unlike previous studies, the pathogen genotypes examined here span a range of relatedness and introduction histories (*i*.*e*., enzootic *vs*. panzootic). Theory predicts that the competitive outcomes among coinfecting pathogens depends on their genetic relatedness (Hamilton 1972, Frank 1996, Brown et al. 2002; Buckling & Brockhurst 2008). Distantly related pathogens are predicted to compete more strongly than closely related (and cooperating) pathogens. However, we found that both facilitation and strong competition occurred among the most distantly related genotypes, P2+E1 *vs*. P1+E1, respectively. Fully testing current theoretical predictions in this system would require a larger and logistically challenging experiment with both intra- and inter-lineage combinations that was beyond the scope of this current study. In the meantime, these results underscore an important point that s often overlooked by current theory: genotype by genotype interactions and introduction history may play stronger roles than relatedness *per se*.

The consistency of virulence patterns in Bd-GPL from both field observations and empirical studies spanning multiple host species is somewhat surprising. Variation within Bd-GPL in phenotypic traits (*e*.*g*., zoosporangia and zoospore size) have been observed in laboratory experiments (Becker et al. 2017; Jenkinson et al. 2018) and field studies (Lambertini et al. 2016), has been linked to pathogen incidence in natural populations (Lambertini et al. 2016) and virulence in experiments (Fisher et al. 2009; Farrer et al. 2011). Additionally, over multiple generations, competitive suppression could select for P1 to invest greater resources into phenotypically-plastic transmission stages such that coinfections could actually lead to higher relative transmission (van Baalen & Sabelis 1995). Within this context, pronounced phenotypic variation in P1 (and other Bd-GPL genotypes) may therefore help explain its ability to successfully outcompete other genotypes, establish a successful infection within-individual hosts and spread through the population, regardless of the high level of virulence. This observed link between competitiveness and virulence in P1 is consistent with theory (van Baalen & Sabelis 1995, Read & Taylor 2001) and empirical studies in several systems (deRoode et al. 2005, Ben-Ami et al. 2008, Inglis et al. 2009, Kinnula et al. 2017, Hodgson et al. 2004, Susi et al. 2015).

Our results agree with previous studies and help to explain why pathogens in general, and Bd in particular, can be highly virulent to their hosts despite their reliance on hosts resources for their own growth and fitness. An important effort for future studies is to understand the extent to which greater phenotypic plasticity in Bd-GPL may enable it to maintain fitness in novel environments and reduce the costs of virulence that are predicted to limit transmission. Identifying the particular genes or pathways that enable such plasticity may offer a novel target for managing outbreaks in Bd and other fungal pathogens.

Mounting evidence indicates that pathogen infections often consist of multiple distinct genotypes (reviewed in Read & Taylor 2001). In the amphibian chytrid, which is implemented in decimating amphibian populations around the globe (Scheele et al. 2019), coinfections with multiple strains may be common and likely to arise largely from the amphibian trade (O’Hanlon et al. 2018; Byrne et al. 2019). We found that coinfections can have important implications for disease dynamics and virulence evolution and warrant further monitoring in wild populations and the pet trade. Thus, understanding how these within-host dynamics play out in the wild over multiple host and pathogen generations carries important implications for the evolutionary trajectory of Bd and any strategies aimed at virulence management in an increasingly globalized society. This underscores the need for ongoing Bd monitoring efforts that include tracking coinfections of multiple Bd genotypes.

## Acknowledgements

We thank Diego Moura Campos, Janaína de Andrade Serrano, Joice Ruggeri Gomes, Mariana Retuci Pontes, Victor Fávaro Augusto, Carolina Lambertini and Kerry Gendreau for technical assistance throughout the experiment. We also thank Skylar Hopkins for comments on the manuscript. This study was approved by the Unicamp Animal Care and Use Committee (CEUA #5398-1/2019), Instituto Chico Mendes de Conservação da Biodiversidade (SISBio #71780-1), and Sistema Nacional de Gestão do Patrimônio Genético e do Conhecimento Tradicional (SISGen #A8246D0). Grants and fellowships were provided by São Paulo Research Foundation (FAPESP #2016/25358-3; #2018/08650-8; #2018/23622-0; #2019/18335-5), the National Council for Scientific and Technological Development (CNPq #300896/2016-6; #302834/2020-6), the Coordination for the Improvement of Higher Education Personnel (CAPES - Finance Code 001), and startup funds from Texas State University to DR and from UW-Madison to JLH.

## Author Contribution

TC conceptualized the study with contributions from CGB and LFT. TC, LPR, DM, DR, and TJ conducted the laboratory methods. TC, DM and JH analyzed the data and drafted the manuscript. All authors carried out the study and critically revised the manuscript.

## Data Availability Statement

The data and full code for all statistical tests are available at dryad (https://insertpostpublication).

