## supplemental_material for "Coinfection with chytrid genotypes drives divergent infection dynamics reflecting broad epidemiological patterns"

[c]Sistema Nacional de Investigación, SENACYT, Building 205, City of Knowledge, Clayton, Panama, Republic of Panama.

[d]Department of Biology, Texas State University, San Marcos, TX 78666, USA.

[e]Department of Biological Sciences, California State University – East Bay, Hayward, CA 94542, USA.

[f]Department of Biology, The Pennsylvania State University, University Park, PA 16802, USA.

[g]School of Veterinary Medicine, Department of Pathobiological Sciences, University of Wisconsin-Madison, WI 53706, USA.

**Keywords:** *Batrachochytrium dendrobatidis*, coinfection, epidemiology, transmission, virulence evolution, within-host interactions

**Table S1.** *Batrachochytrium dendrobatidis* (Bd) genotypes isolated from amphibians across Brazilian Atlantic Forest. This table includes isolate name or museum record of the host, host species, sampled host life-stage (tadpole or adult) and breeding habitat, collector name, isolation year, chytrid lineage or genotype, municipality, state, latitude (Lat), longitude (Lon), and reference.

| Isolate/Musem record | Host species | Habitat | Stage | Collector | Isolation year | Lineage | Municipality | State | Lat | Lon | Reference |
|---|---|---|---|---|---|---|---|---|---|---|---|
| CLFT 001/10 | *Hylodes japi* | Stream | Tadpole | CA Vieira | 2010 | Bd-Asia-2/ Brazil | Jundiaí | SP | -23.25 | -46.95 | CLFT |
| CLFT 021 | Unidentified | Unknown | Tadpole | CA Vieira | 2010 | Bd-GPL | Cabreúva | SP | -23.31 | -47.10 | CLFT |
| CLFT 023 | *Boana* sp. | Pond | Tadpole | CA Vieira | 2011 | Bd-GPL | Camanducaia | MG | -22.87 | -46.03 | CLFT |
| CLFT 024-01 | *Hylodes cardosoi* | Stream | Tadpole | CA Vieira | 2011 | Bd-GPL | Morretes | PR | -25.36 | -48.88 | CLFT |
| CLFT 024-02 | *Hylodes cardosoi* | Stream | Tadpole | CA Vieira | 2011 | Hybrid | Morretes | PR | -25.36 | -48.88 | CLFT |
| CLFT 026 | *Boana faber* | Pond | Tadpole | C Lambertini | 2011 | Bd-GPL | Iporanga | SP | -24.58 | -48.60 | CLFT |
| CLFT 029-00 | *Boana* cf. *albopunctata* | Pond | Tadpole | C Lambertini | 2011 | Bd-GPL | Jundiaí | SP | -23.25 | -46.95 | CLFT |
| CLFT 029-01 | *Scinax hiemalis* | Pond | Tadpole | C Lambertini | 2011 | Bd-GPL | Jundiaí | SP | -23.25 | -46.95 | CLFT |
| CLFT 030 | *Hylodes phyllodes* | Stream | Tadpole | C Lambertini | 2012 | Bd-GPL | Bertioga | SP | -23.71 | -46.03 | CLFT |
| CLFT 031 | *Hylodes phyllodes* | Stream | Tadpole | C Lambertini | 2012 | Bd-GPL | Bertioga | SP | -23.71 | -46.03 | CLFT |
| CLFT 032 | *Hylodes phyllodes* | Stream | Tadpole | C Lambertini | 2012 | Bd-GPL | Bertioga | SP | -23.71 | -46.03 | CLFT |
| CLFT 033 | *Hylodes phyllodes* | Stream | Tadpole | C Lambertini | 2012 | Bd-GPL | Bertioga | SP | -23.71 | -46.03 | CLFT |
| CLFT 034 | *Hylodes phyllodes* | Stream | Tadpole | TS Jenkinson | 2013 | Bd-GPL | Bertioga | SP | -23.71 | -46.03 | CLFT |
| CLFT 035 | *Boana faber* | Pond | Tadpole | KR Zamudio | 2013 | Bd-GPL | Iporanga | SP | -24.58 | -48.60 | CLFT |
| CLFT 036 | *Boana faber* | Pond | Tadpole | D Rodriguez | 2013 | Bd-GPL | Iporanga | SP | -24.58 | -48.60 | CLFT |

| | | | | | | | | | | | |
|---|---|---|---|---|---|---|---|---|---|---|---|
| CLFT 037 | *Boana faber* | Pond | Tadpole | KR Zamudio | 2013 | Bd-GPL | Iporanga | SP | -24.58 | -48.60 | CLFT |
| CLFT 038 | *Bokermannohyla hylax* | Stream | Tadpole | TS Jenkinson | 2013 | Hybrid | Morretes | PR | -25.36 | -48.88 | CLFT |
| CLFT 039 | *Bokermannohyla hylax* | Stream | Tadpole | TS Jenkinson | 2013 | Hybrid | Morretes | PR | -25.36 | -48.88 | CLFT |
| CLFT 040 | *Bokermannohyla hylax* | Stream | Tadpole | LF Toledo | 2013 | Bd-Asia-2/ Brazil | Morretes | PR | -25.36 | -48.88 | CLFT |
| CLFT 041 | *Bokermannohyla hylax* | Stream | Tadpole | D Rodriguez | 2013 | Bd-Asia-2/ Brazil | Morretes | PR | -25.36 | -48.88 | CLFT |
| CLFT 042 | *Boana faber* | Pond | Tadpole | C Betancourt | 2013 | Bd-GPL | Iporanga | SP | -24.58 | -48.60 | CLFT |
| CLFT 043 | *Bokermannohyla hylax* | Stream | Tadpole | TS Jenkinson | 2013 | Bd-GPL | Morretes | PR | -25.36 | -48.88 | CLFT |
| CLFT 044 | *Hylodes cardosoi* | Stream | Tadpole | C Betancourt | 2013 | Bd-Asia-2/ Brazil | Morretes | PR | -25.36 | -48.88 | CLFT |
| CLFT 045 | *Hylodes cardosoi* | Stream | Tadpole | TS Jenkinson | 2013 | Bd-GPL | Morretes | PR | -25.36 | -48.88 | CLFT |
| CLFT 046 | *Bokermannohyla hylax* | Stream | Tadpole | C Betancourt | 2013 | Bd-GPL | Morretes | PR | -25.36 | -48.88 | CLFT |
| CLFT 047 | *Bokermannohyla hylax* | Stream | Tadpole | C Betancourt | 2013 | Bd-GPL | Morretes | PR | -25.36 | -48.88 | CLFT |
| CLFT 048 | *Hylodes meridionalis* | Stream | Tadpole | C Betancourt | 2013 | Bd-GPL | Rancho Queimado | SC | -27.67 | -49.09 | CLFT |
| CLFT 049 | *Hylodes meridionalis* | Stream | Tadpole | TS Jenkinson | 2013 | Bd-GPL | Rancho Queimado | SC | -27.67 | -49.09 | CLFT |
| CLFT 050 | *Hylodes meridionalis* | Stream | Tadpole | C Betancourt | 2013 | Bd-GPL | Rancho Queimado | SC | -27.67 | -49.09 | CLFT |
| CLFT 051 | *Hylodes meridionalis* | Stream | Tadpole | TS Jenkinson | 2013 | Bd-GPL | Rancho Queimado | SC | -27.67 | -49.09 | CLFT |
| CLFT 052 | *Hylodes meridionalis* | Stream | Tadpole | C Betancourt | 2013 | Bd-GPL | Rancho Queimado | SC | -27.67 | -49.09 | CLFT |

| | | | | | | | | | | | |
|---|---|---|---|---|---|---|---|---|---|---|---|
| CLFT 053 | *Hylodes meridionalis* | Stream | Tadpole | KR Zamudio | 2013 | Bd-GPL | Rancho Queimado | SC | -27.67 | -49.09 | CLFT |
| CLFT 054 | *Hylodes meridionalis* | Stream | Tadpole | D Rodriguez | 2013 | Bd-GPL | Rancho Queimado | SC | -27.67 | -49.09 | CLFT |
| CLFT 055 | *Hylodes meridionalis* | Stream | Tadpole | TY James | 2013 | Bd-GPL | Rancho Queimado | SC | -27.67 | -49.09 | CLFT |
| CLFT 056 | *Hylodes meridionalis* | Stream | Tadpole | TS Jenkinson | 2013 | Bd-GPL | Rancho Queimado | SC | -27.67 | -49.09 | CLFT |
| CLFT 057 | *Hylodes meridionalis* | Stream | Tadpole | C Betancourt | 2013 | Bd-GPL | Rancho Queimado | SC | -27.67 | -49.09 | CLFT |
| CLFT 058 | *Hylodes meridionalis* | Stream | Tadpole | TS Jenkinson | 2013 | Bd-GPL | Rancho Queimado | SC | -27.67 | -49.09 | CLFT |
| CLFT 060 | *Hylodes meridionalis* | Stream | Tadpole | TS Jenkinson | 2013 | Bd-GPL | Pomerode | SC | -26.77 | -49.19 | CLFT |
| CLFT 061 | *Hylodes meridionalis* | Stream | Tadpole | C Betancourt | 2013 | Bd-Asia-2/ Brazil | Pomerode | SC | -26.77 | -49.19 | CLFT |
| CLFT 062 | *Hylodes meridionalis* | Stream | Tadpole | C Betancourt | 2013 | Bd-GPL | Pomerode | SC | -26.77 | -49.19 | CLFT |
| CLFT 063 | *Hylodes meridionalis* | Stream | Tadpole | C Betancourt | 2013 | Bd-GPL | Pomerode | SC | -26.77 | -49.19 | CLFT |
| CLFT 064 | *Hylodes meridionalis* | Stream | Tadpole | C Betancourt | 2013 | Bd-GPL | Pomerode | SC | -26.77 | -49.19 | CLFT |
| CLFT 065 | *Hylodes japi* | Stream | Tadpole | C Betancourt | 2013 | Bd-Asia-2/ Brazil | Jundiaí | SP | -23.25 | -46.95 | CLFT |
| CLFT 066 | *Hylodes japi* | Stream | Tadpole | J Longcore | 2013 | Bd-Asia-2/ Brazil | Jundiaí | SP | -23.25 | -46.95 | CLFT |
| CLFT 067 | *Hylodes japi* | Stream | Tadpole | C Betancourt | 2013 | Bd-Asia-2/ Brazil | Jundiaí | SP | -23.25 | -46.95 | CLFT |
| CLFT 068 | *Hylodes japi* | Stream | Tadpole | C Betancourt | 2013 | Bd-Asia-2/ Brazil | Jundiaí | SP | -23.25 | -46.95 | CLFT |
| CLFT 070 | *Hylodes japi* | Stream | Tadpole | J Longcore | 2013 | Bd-Asia-2/ Brazil | Jundiaí | SP | -23.25 | -46.95 | CLFT |

| | | | | | | | | | | | |
|---|---|---|---|---|---|---|---|---|---|---|---|
| CLFT 071 | *Hylodes japi* | Stream | Tadpole | C Betancourt | 2013 | Bd-Asia-2/ Brazil | Jundiaí | SP | -23.25 | -46.95 | CLFT |
| CLFT 073 | *Aplastodiscus* sp. | Stream | Tadpole | C Betancourt | 2013 | Bd-GPL | Teresópolis | RJ | -22.46 | -43.00 | CLFT |
| CLFT 074 | Unidentified | Stream | Tadpole | C Betancourt | 2013 | Bd-GPL | Teresópolis | RJ | -22.46 | -43.00 | CLFT |
| CLFT 075 | Unidentified | Stream | Tadpole | TY James | 2013 | Bd-GPL | Teresópolis | RJ | -22.46 | -43.00 | CLFT |
| CLFT 076 | *Bokermannohyla* sp. | Stream | Tadpole | C Betancourt | 2013 | Bd-GPL | Teresópolis | RJ | -22.46 | -43.00 | CLFT |
| CLFT 077 | *Bokermannohyla* sp. | Stream | Tadpole | C Betancourt | 2013 | Bd-GPL | Teresópolis | RJ | -22.46 | -43.00 | CLFT |
| CLFT 078 | *Bokermannohyla* sp. | Stream | Tadpole | TY James | 2013 | Bd-GPL | Teresópolis | RJ | -22.46 | -43.00 | CLFT |
| CLFT 079 | *Bokermannohyla* sp. | Stream | Tadpole | TY James | 2013 | Bd-GPL | Teresópolis | RJ | -22.46 | -43.00 | CLFT |
| CLFT 080 | *Bokermannohyla* sp. | Stream | Tadpole | C Betancourt | 2013 | Bd-GPL | Teresópolis | RJ | -22.46 | -43.00 | CLFT |
| CLFT 081 | Unidentified | Stream | Tadpole | C Betancourt | 2013 | Bd-GPL | Teresópolis | RJ | -22.46 | -43.00 | CLFT |
| CLFT 082 | *Bokermannohyla* sp. | Stream | Tadpole | C Betancourt | 2013 | Bd-GPL | Teresópolis | RJ | -22.46 | -43.00 | CLFT |
| CLFT 083 | *Scinax hayii* | Pond | Tadpole | C Betancourt | 2013 | Bd-GPL | Teresópolis | RJ | -22.46 | -43.00 | CLFT |
| CLFT 084 | *Bokermannohyla* sp. | Stream | Tadpole | C Betancourt | 2013 | Bd-GPL | Teresópolis | RJ | -22.46 | -43.00 | CLFT |
| CLFT 085 | Unidentified | Stream | Tadpole | C Betancourt | 2013 | Bd-GPL | Teresópolis | RJ | -22.46 | -43.00 | CLFT |
| CLFT 086 | Unidentified | Stream | Tadpole | C Betancourt | 2013 | Bd-GPL | Teresópolis | RJ | -22.46 | -43.00 | CLFT |
| CLFT 087 | *Scinax hayii* | Pond | Tadpole | C Betancourt | 2013 | Bd-GPL | Teresópolis | RJ | -22.46 | -43.00 | CLFT |
| CLFT 088 | *Scinax hayii* | Pond | Tadpole | C Betancourt & TS Jenkison | 2013 | Bd-GPL | Teresópolis | RJ | -22.46 | -43.00 | CLFT |
| CLFT 089 | *Aplastodiscus sibilatus* | Stream/ Pond | Tadpole | LFM Lima | 2013 | Bd-GPL | Maceió | AL | -9.23 | -35.86 | CLFT |

| | | | | | | | | | | | |
|---|---|---|---|---|---|---|---|---|---|---|---|
| CLFT 090 | *Aplastodiscus sibilatus* | Stream/ Pond | Tadpole | LFM Lima | 2013 | Bd-GPL | Maceió | AL | -9.23 | -35.86 | CLFT |
| CLFT 091 | *Aplastodiscus sibilatus* | Stream/ Pond | Tadpole | LFM Lima | 2013 | Bd-GPL | Maceió | AL | -9.23 | -35.86 | CLFT |
| CLFT 092 | *Aplastodiscus sibilatus* | Stream/ Pond | Tadpole | ABCR Lima | 2013 | Bd-GPL | Maceió | AL | -9.23 | -35.86 | CLFT |
| CLFT 093 | *Aplastodiscus sibilatus* | Stream/ Pond | Tadpole | ABCR Lima | 2013 | Bd-GPL | Maceió | AL | -9.23 | -35.86 | CLFT |
| CLFT 094 | *Aplastodiscus sibilatus* | Stream/ Pond | Tadpole | C Lambertini | 2013 | Bd-GPL | Maceió | AL | -9.23 | -35.86 | CLFT |
| CLFT 095 | *Aplastodiscus* sp. | Stream | Tadpole | TS Jenkinson | 2014 | Bd-GPL | Camacan | BA | -15.38 | -39.57 | CLFT |
| CLFT 096 | *Aplastodiscus* sp. | Stream | Tadpole | C Lambertini | 2014 | Bd-GPL | Camacan | BA | -15.38 | -39.57 | CLFT |
| CLFT 097 | *Aplastodiscus* sp. | Stream | Tadpole | AV Aguilar | 2014 | Bd-GPL | Camacan | BA | -15.38 | -39.57 | CLFT |
| CLFT 098 | *Aplastodiscus* sp. | Stream | Tadpole | C Lambertini | 2014 | Bd-GPL | Camacan | BA | -15.38 | -39.57 | CLFT |
| CLFT 099 | *Aplastodiscus* sp. | Stream | Tadpole | TS Jenkinson | 2014 | Bd-GPL | Camacan | BA | -15.38 | -39.57 | CLFT |
| CLFT 100 | *Bokermannohyla* sp. | Stream | Tadpole | C Lambertini | 2014 | Bd-GPL | Camacan | BA | -15.38 | -39.57 | CLFT |
| CLFT 101 | *Aplastodiscus* sp. | Stream | Tadpole | TS Jenkinson | 2014 | Bd-GPL | Camacan | BA | -15.38 | -39.57 | CLFT |
| CLFT 102 | *Bokermannohyla* sp. | Stream | Tadpole | TS Jenkinson | 2014 | Bd-GPL | Camacan | BA | -15.38 | -39.57 | CLFT |
| CLFT 103 | *Bokermannohyla* sp. | Stream | Tadpole | AV Aguilar | 2014 | Bd-GPL | Camacan | BA | -15.38 | -39.57 | CLFT |
| CLFT 104 | *Bokermannohyla* sp. | Stream | Tadpole | AV Aguilar | 2014 | Bd-GPL | Camacan | BA | -15.38 | -39.57 | CLFT |
| CLFT 105 | *Bokermannohyla* sp. | Stream | Tadpole | TS Jenkinson | 2014 | Bd-GPL | Camacan | BA | -15.38 | -39.57 | CLFT |
| CLFT 106 | *Bokermannohyla* sp. | Stream | Tadpole | TS Jenkinson | 2014 | Bd-GPL | Camacan | BA | -15.38 | -39.57 | CLFT |

| | | | | | | | | | | | |
|---|---|---|---|---|---|---|---|---|---|---|---|
| CLFT 107 | *Bokermannohyla* sp. | Stream | Tadpole | C Lambertini | 2014 | Bd-GPL | Camacan | BA | -15.38 | -39.57 | CLFT |
| CLFT 108 | *Bokermannohyla* sp. | Stream | Tadpole | TS Jenkinson | 2014 | Bd-GPL | Camacan | BA | -15.38 | -39.57 | CLFT |
| CLFT 109 | *Bokermannohyla* sp. | Stream | Tadpole | TS Jenkinson | 2014 | Bd-GPL | Camacan | BA | -15.38 | -39.57 | CLFT |
| CLFT 110 | *Bokermannohyla* sp. | Stream | Tadpole | AV Aguilar | 2014 | Bd-GPL | Camacan | BA | -15.38 | -39.57 | CLFT |
| CLFT 111 | *Bokermannohyla* sp. | Stream | Tadpole | TS Jenkinson | 2014 | Bd-GPL | Santa Teresa | ES | -19.83 | -40.59 | CLFT |
| CLFT 112 | *Aplastodiscus* sp. | Stream | Tadpole | AV Aguilar | 2014 | Bd-GPL | Santa Teresa | ES | -19.83 | -40.59 | CLFT |
| CLFT 113 | Unidentified | Stream | Tadpole | TS Jenkinson | 2014 | Bd-GPL | Santa Teresa | ES | -19.83 | -40.59 | CLFT |
| CLFT 114 | *Bokermannohyla* sp. | Stream | Tadpole | AV Aguilar | 2014 | Bd-GPL | Santa Teresa | ES | -19.83 | -40.59 | CLFT |
| CLFT 115 | *Bokermannohyla* sp. | Stream | Tadpole | AV Aguilar | 2014 | Bd-GPL | Santa Teresa | ES | -19.83 | -40.59 | CLFT |
| CLFT 116 | *Bokermannohyla* sp. | Stream | Tadpole | TS Jenkinson | 2014 | Bd-GPL | Santa Teresa | ES | -19.83 | -40.59 | CLFT |
| CLFT 117 | *Bokermannohyla* sp. | Stream | Tadpole | C Lambertini | 2014 | Bd-GPL | Santa Teresa | ES | -19.83 | -40.59 | CLFT |
| CLFT 118 | *Bokermannohyla* sp. | Stream | Tadpole | AV Aguilar | 2014 | Bd-GPL | Santa Teresa | ES | -19.83 | -40.59 | CLFT |
| CLFT 119 | *Bokermannohyla* sp. | Stream | Tadpole | AV Aguilar | 2014 | Bd-GPL | Santa Teresa | ES | -19.83 | -40.59 | CLFT |
| CLFT 120 | *Bokermannohyla* sp. | Stream | Tadpole | AV Aguilar | 2014 | Bd-GPL | Santa Teresa | ES | -19.83 | -40.59 | CLFT |
| CLFT 121 | *Bokermannohyla* sp. | Stream | Tadpole | AV Aguilar | 2014 | Bd-GPL | Santa Teresa | ES | -19.83 | -40.59 | CLFT |
| CLFT 122 | *Bokermannohyla* sp. | Stream | Tadpole | TS Jenkinson | 2014 | Bd-GPL | Santa Teresa | ES | -19.83 | -40.59 | CLFT |
| CLFT 123 | *Bokermannohyla* sp. | Stream | Tadpole | C Lambertini | 2014 | Bd-GPL | Santa Teresa | ES | -19.83 | -40.59 | CLFT |
| CLFT 124 | *Bokermannohyla* sp. | Stream | Tadpole | AV Aguilar | 2014 | Bd-GPL | Santa Teresa | ES | -19.83 | -40.59 | CLFT |
| CLFT 125 | Unidentified | Stream | Tadpole | C Lambertini | 2014 | Bd-GPL | Santa Teresa | ES | -19.83 | -40.59 | CLFT |
| CLFT 126 | *Phyllomedusa* sp. | Pond | Tadpole | AV Aguilar | 2014 | Bd-GPL | Vargem Alta | ES | -20.66 | -41.00 | CLFT |

| | | | | | | | | | | | |
|---|---|---|---|---|---|---|---|---|---|---|---|
| CLFT 127 | *Dendropsophus minutus* | Pond | Tadpole | TY James | 2014 | Bd-GPL | Vargem Alta | ES | -20.66 | -41.00 | CLFT |
| CLFT 128 | *Aplastodiscus* sp. | Stream | Tadpole | TY James | 2014 | Bd-GPL | Vargem Alta | ES | -20.66 | -41.00 | CLFT |
| CLFT 129 | *Aplastodiscus* sp. | Stream | Tadpole | AV Aguilar | 2014 | Bd-GPL | Vargem Alta | ES | -20.66 | -41.00 | CLFT |
| CLFT 130 | *Scinax fuscovarius* | Stream | Tadpole | AV Aguilar | 2014 | Bd-GPL | Vargem Alta | ES | -20.66 | -41.00 | CLFT |
| CLFT 131 | *Aquarana catesbeiana* | Pond | Tadpole | TS Jenkinson | 2014 | Bd-GPL | Vargem Alta | ES | -20.66 | -41.00 | CLFT |
| CLFT 132 | *Dendropsophus minutus* | Pond | Tadpole | AV Aguilar | 2014 | Bd-GPL | Vargem Alta | ES | -20.66 | -41.00 | CLFT |
| CLFT 133 | *Phyllomedusa* sp. | Pond | Tadpole | AV Aguilar | 2014 | Bd-GPL | Vargem Alta | ES | -20.66 | -41.00 | CLFT |
| CLFT 134 | *Phyllomedusa* sp. | Pond | Tadpole | TS Jenkinson | 2014 | Bd-GPL | Vargem Alta | ES | -20.66 | -41.00 | CLFT |
| CLFT 135 | *Scinax fuscovarius* | Pond | Tadpole | KR Zamudio | 2014 | Bd-GPL | Vargem Alta | ES | -20.66 | -41.00 | CLFT |
| CLFT 136 | *Bokermannohyla hylax* | Stream | Tadpole | TS Jenkinson | 2014 | Bd-Asia-2/ Brazil | Morretes | PR | -25.36 | -48.88 | CLFT |
| CLFT 137 | *Hylodes cardosoi* | Stream | Tadpole | TS Jenkinson | 2014 | Bd-GPL | Morretes | PR | -25.36 | -48.88 | CLFT |
| CLFT 138 | *Hylodes cardosoi* | Stream | Tadpole | C Lambertini | 2014 | Bd-GPL | Morretes | PR | -25.36 | -48.88 | CLFT |
| CLFT 139 | *Hylodes cardosoi* | Stream | Tadpole | TS Jenkinson | 2014 | Bd-Asia-2/ Brazil | Morretes | PR | -25.36 | -48.88 | CLFT |
| CLFT 141 | *Hylodes cardosoi* | Stream | Tadpole | LFM Lima | 2014 | Bd-Asia-2/ Brazil | Morretes | PR | -25.36 | -48.88 | CLFT |
| CLFT 142 | *Crossodactylus caramaschii* | Stream | Tadpole | PP Morao | 2014 | Bd-Asia-2/ Brazil | Morretes | PR | -25.36 | -48.88 | CLFT |
| CLFT 143 | *Hylodes cardosoi* | Stream | Tadpole | TS Jenkinson | 2014 | Bd-Asia-2/ Brazil | Morretes | PR | -25.36 | -48.88 | CLFT |

| | | | | | | | | | | | |
|---|---|---|---|---|---|---|---|---|---|---|---|
| CLFT 144 | *Hylodes cardosoi* | Stream | Tadpole | TS Jenkinson | 2014 | Bd-Asia-2/ Brazil | Morretes | PR | -25.36 | -48.88 | CLFT |
| CLFT 145 | *Hylodes cardosoi* | Stream | Tadpole | PP Morao | 2014 | Bd-Asia-2/ Brazil | Morretes | PR | -25.36 | -48.88 | CLFT |
| CLFT 146 | *Hylodes cardosoi* | Stream | Tadpole | TS Jenkinson | 2014 | Bd-Asia-2/ Brazil | Morretes | PR | -25.36 | -48.88 | CLFT |
| CLFT 148 | *Hylodes cardosoi* | Stream | Tadpole | TS Jenkinson | 2014 | Bd-Asia-2/ Brazil | Morretes | PR | -25.36 | -48.88 | CLFT |
| CLFT 149 | *Hylodes cardosoi* | Stream | Tadpole | TS Jenkinson | 2014 | Bd-Asia-2/ Brazil | Morretes | PR | -25.36 | -48.88 | CLFT |
| CLFT 150 | *Hylodes cardosoi* | Stream | Tadpole | PP Morao | 2014 | Bd-Asia-2/ Brazil | Morretes | PR | -25.36 | -48.88 | CLFT |
| CLFT 151 | *Hylodes cardosoi* | Stream | Tadpole | PP Morao | 2014 | Bd-Asia-2/ Brazil | Morretes | PR | -25.36 | -48.88 | CLFT |
| CLFT 152 | *Crossodactylus caramaschii* | Stream | Tadpole | TS Jenkinson | 2014 | Bd-GPL | Morretes | PR | -25.36 | -48.88 | CLFT |
| CLFT 153 | *Hylodes cardosoi* | Stream | Tadpole | TS Jenkinson | 2014 | Bd-Asia-2/ Brazil | Morretes | PR | -25.36 | -48.88 | CLFT |
| CLFT 154 | *Hylodes cardosoi* | Stream | Tadpole | TS Jenkinson | 2014 | Bd-GPL | Morretes | PR | -25.36 | -48.88 | CLFT |
| CLFT 156 | *Hylodes cardosoi* | Stream | Tadpole | TY James | 2015 | Bd-GPL | Morretes | PR | -25.36 | -48.88 | CLFT |
| CLFT 157 | *Hylodes cardosoi* | Stream | Tadpole | TS Jenkinson | 2015 | Bd-GPL | Morretes | PR | -25.36 | -48.88 | CLFT |
| CLFT 158 | *Hylodes cardosoi* | Stream | Tadpole | TY James | 2015 | Bd-GPL | Morretes | PR | -25.36 | -48.88 | CLFT |
| CLFT 159 | *Hylodes cardosoi* | Stream | Tadpole | T Carvalho | 2015 | Bd-GPL | Morretes | PR | -25.57 | -48.90 | CLFT |
| CLFT 160 | *Hylodes cardosoi* | Stream | Tadpole | TS Jenkinson | 2015 | Hybrid | Morretes | PR | -25.57 | -48.90 | CLFT |

| | | | | | | | | | | | |
|---|---|---|---|---|---|---|---|---|---|---|---|
| CLFT 161 | *Hylodes cardosoi* | Stream | Tadpole | TY James | 2015 | Bd-GPL | Morretes | PR | -25.57 | -48.90 | CLFT |
| CLFT 162 | *Hylodes cardosoi* | Stream | Tadpole | TS Jenkinson | 2015 | Bd-GPL | Morretes | PR | -25.57 | -48.90 | CLFT |
| CLFT 163 | *Hylodes cardosoi* | Stream | Tadpole | TS Jenkinson | 2015 | Bd-GPL | Morretes | PR | -25.57 | -48.90 | CLFT |
| CLFT 164 | *Hylodes cardosoi* | Stream | Tadpole | TY James | 2015 | Bd-GPL | Morretes | PR | -25.57 | -48.90 | CLFT |
| CLFT 165 | *Hylodes cardosoi* | Stream | Tadpole | TS Jenkinson | 2015 | Hybrid | Morretes | PR | -25.36 | -48.88 | CLFT |
| CLFT 166 | Unidentified | Stream | Tadpole | PP Morao | 2015 | Bd-GPL | Campos do Jordão | SP | -22.78 | -45.62 | CLFT |
| CLFT 167 | Unidentified | Stream | Tadpole | TS Jenkinson | 2015 | Bd-GPL | Campos do Jordão | SP | -22.78 | -45.62 | CLFT |
| CLFT 168 | *Phantasmarana apuana* | Stream | Tadpole | C Lambertini | 2015 | Bd-GPL | Alto Caparaó | MG | -20.42 | -41.85 | CLFT |
| CLFT 169 | *Phantasmarana apuana* | Stream | Tadpole | LFM de Lima | 2015 | Bd-GPL | Alto Caparaó | MG | -20.42 | -41.85 | CLFT |
| CLFT 170 | *Aquarana catesbeiana* | Farm | Tadpole | LP Ribeiro | 2016 | Bd-Asia-2/ Brazil | Pindamonhangaba | SP | -22.85 | -45.49 | CLFT |
| CLFT 171 | *Aquarana catesbeiana* | Farm | Tadpole | LP Ribeiro | 2016 | Bd-Asia-2/ Brazil | Pindamonhangaba | SP | -22.85 | -45.49 | CLFT |
| CLFT 172 | *Aquarana catesbeiana* | Farm | Tadpole | LP Ribeiro | 2016 | Bd-Asia-2/ Brazil | Pindamonhangaba | SP | -22.85 | -45.49 | CLFT |
| CLFT 173 | *Aquarana catesbeiana* | Farm | Tadpole | LP Ribeiro | 2016 | Bd-GPL | Pindamonhangaba | SP | -22.85 | -45.49 | CLFT |
| CLFT 174 | *Aquarana catesbeiana* | Farm | Tadpole | LP Ribeiro | 2016 | Bd-GPL | Pindamonhangaba | SP | -22.85 | -45.49 | CLFT |
| CLFT 175 | *Aquarana catesbeiana* | Farm | Tadpole | LP Ribeiro | 2016 | Bd-Asia-2/ Brazil | Pindamonhangaba | SP | -22.85 | -45.49 | CLFT |

| | | | | | | | | | | | |
|---|---|---|---|---|---|---|---|---|---|---|---|
| CLFT 176 | *Aquarana catesbeiana* | Farm | Tadpole | LP Ribeiro | 2016 | Bd-GPL | Pindamonhangaba | SP | -22.85 | -45.49 | CLFT |
| CLFT 177 | *Aquarana catesbeiana* | Farm | Tadpole | LP Ribeiro | 2016 | Bd-GPL | Pindamonhangaba | SP | -22.85 | -45.49 | CLFT |
| CLFT 178 | *Aquarana catesbeiana* | Farm | Tadpole | LP Ribeiro | 2016 | Bd-GPL | Pindamonhangaba | SP | -22.85 | -45.49 | CLFT |
| CLFT 179 | *Aquarana catesbeiana* | Farm | Tadpole | LP Ribeiro | 2016 | Bd-GPL | Pindamonhangaba | SP | -22.85 | -45.49 | CLFT |
| CLFT 180 | *Aquarana catesbeiana* | Farm | Tadpole | LP Ribeiro | 2016 | Bd-GPL | Pindamonhangaba | SP | -22.85 | -45.49 | CLFT |
| CLFT 181 | *Aquarana catesbeiana* | Farm | Tadpole | LP Ribeiro | 2016 | Bd-GPL | Pindamonhangaba | SP | -22.85 | -45.49 | CLFT |
| CLFT 182 | *Aquarana catesbeiana* | Farm | Tadpole | LP Ribeiro | 2016 | Bd-GPL | Pindamonhangaba | SP | -22.85 | -45.49 | CLFT |
| CLFT 183 | *Aquarana catesbeiana* | Farm | Tadpole | LP Ribeiro | 2016 | Bd-Asia-2/ Brazil | Pindamonhangaba | SP | -22.85 | -45.49 | CLFT |
| CLFT 184 | *Aquarana catesbeiana* | Farm | Tadpole | LP Ribeiro | 2016 | Bd-GPL | Santa Barbara D’Oeste | SP | -22.76 | -47.41 | CLFT |
| CLFT 185 | *Aquarana catesbeiana* | Farm | Tadpole | LP Ribeiro | 2016 | Bd-GPL | Santa Barbara D’Oeste | SP | -22.76 | -47.41 | CLFT |
| CLFT 186 | *Aquarana catesbeiana* | Farm | Tadpole | LP Ribeiro | 2016 | Bd-GPL | Santa Barbara D’Oeste | SP | -22.76 | -47.41 | CLFT |
| CLFT 187 | *Aquarana catesbeiana* | Farm | Tadpole | LP Ribeiro | 2016 | Bd-GPL | Santa Barbara D’Oeste | SP | -22.76 | -47.41 | CLFT |
| CLFT 188 | *Aquarana catesbeiana* | Farm | Tadpole | LP Ribeiro | 2016 | Bd-GPL | Santa Barbara D’Oeste | SP | -22.76 | -47.41 | CLFT |

| | | | | | | | | | | | |
|---|---|---|---|---|---|---|---|---|---|---|---|
| CLFT 189 | *Aquarana catesbeiana* | Farm | Tadpole | LP Ribeiro | 2016 | Bd-GPL | Santa Barbara D’Oeste | SP | -22.76 | -47.41 | CLFT |
| CLFT 190 | *Aquarana catesbeiana* | Farm | Tadpole | LP Ribeiro | 2016 | Bd-GPL | Santa Barbara D’Oeste | SP | -22.76 | -47.41 | CLFT |
| CLFT 191 | *Aquarana catesbeiana* | Farm | Tadpole | LP Ribeiro | 2016 | Bd-GPL | Santa Isabel | SP | -23.32 | -46.23 | CLFT |
| CLFT 192 | *Aquarana catesbeiana* | Farm | Tadpole | LP Ribeiro | 2016 | Bd-GPL | Santa Isabel | SP | -23.32 | -46.23 | CLFT |
| CLFT 193 | *Aquarana catesbeiana* | Farm | Tadpole | LP Ribeiro | 2016 | Bd-GPL | Santa Isabel | SP | -23.32 | -46.23 | CLFT |
| CLFT 194 | *Aquarana catesbeiana* | Farm | Tadpole | LP Ribeiro | 2016 | Bd-GPL | Santa Isabel | SP | -23.32 | -46.23 | CLFT |
| CLFT 195 | *Aquarana catesbeiana* | Farm | Tadpole | LP Ribeiro | 2016 | Bd-GPL | Santa Isabel | SP | -23.32 | -46.23 | CLFT |
| CLFT 196 | *Aquarana catesbeiana* | Farm | Tadpole | LP Ribeiro | 2016 | Bd-GPL | Santa Isabel | SP | -23.32 | -46.23 | CLFT |
| CLFT 197 | *Aquarana catesbeiana* | Farm | Tadpole | LP Ribeiro | 2016 | Bd-GPL | Santa Isabel | SP | -23.32 | -46.23 | CLFT |
| CLFT 198 | *Aquarana catesbeiana* | Farm | Tadpole | LP Ribeiro | 2016 | Bd-GPL | Santa Isabel | SP | -23.32 | -46.23 | CLFT |
| CLFT 199 | Unidentified | Unknown | Tadpole | LP Ribeiro & T Carvalho | 2016 | Bd-GPL | Jundiaí | SP | -23.25 | -46.95 | CLFT |
| CLFT 201 | Unidentified | Unknown | Tadpole | C Lambertini | 2016 | Bd-GPL | Capitólio | MG | -20.65 | -46.26 | CLFT |
| CLFT 202 | *Aquarana catesbeiana* | Farm | Tadpole | LP Ribeiro | 2016 | Bd-Asia-2/ Brazil | São Paulo | SP | -23.55 | -46.63 | CLFT |

| | | | | | | | | | | | |
|---|---|---|---|---|---|---|---|---|---|---|---|
| CLFT 203 | *Aquarana catesbeiana* | Farm | Tadpole | LP Ribeiro | 2016 | Bd-Asia-2/ Brazil | São Paulo | SP | -23.55 | -46.63 | CLFT |
| CLFT 204 | *Aquarana catesbeiana* | Farm | Tadpole | LP Ribeiro | 2016 | Bd-Asia-2/ Brazil | São Paulo | SP | -23.55 | -46.63 | CLFT |
| CLFT 205 | *Aquarana catesbeiana* | Farm | Tadpole | LP Ribeiro | 2016 | Bd-Asia-2/ Brazil | São Paulo | SP | -23.55 | -46.63 | CLFT |
| CLFT 206 | *Aquarana catesbeiana* | Farm | Tadpole | LP Ribeiro | 2016 | Bd-GPL | São Paulo | SP | -23.55 | -46.63 | CLFT |
| CLFT 207 | *Aquarana catesbeiana* | Farm | Tadpole | LP Ribeiro | 2016 | Bd-Asia-2/ Brazil | São Paulo | SP | -23.55 | -46.63 | CLFT |
| CLFT 209 | *Aquarana catesbeiana* | Farm | Tadpole | LP Ribeiro | 2016 | Bd-Asia-2/ Brazil | São Paulo | SP | -23.55 | -46.63 | CLFT |
| JEL 648 | *Hylodes japi* | Stream | Tadpole | J Longcore | 2010 | Bd-Asia-2/ Brazil | Jundiaí | SP | -23.25 | -46.95 | Schloegel et al. 2012 |
| JEL 649 | *Hylodes japi* | Stream | Tadpole | J Longcore | 2010 | Bd-Asia-2/ Brazil | Jundiaí | SP | -23.25 | -46.95 | Schloegel et al. 2012 |
| LMS 902 | *Aquarana catesbeiana* | Farm | Adult | LM Schloegel | 2008 | Bd-GPL | Pindamonhangaba | SP | -22.85 | -45.49 | Schloegel et al. 2012 |
| LMS 925 | *Aquarana catesbeiana* | Farm | Adult | LM Schloegel | 2008 | Bd-GPL | Pindamonhangaba | SP | -22.85 | -45.49 | Schloegel et al. 2012 |
| LMS 931 | *Aquarana catesbeiana* | Farm | Adult | LM Schloegel | 2009 | Bd-GPL | Tremembé | SP | -23.00 | -45.51 | Schloegel et al. 2012 |
| MNRJ 1352 | *Cycloramphus brasiliensis* | Stream | Adult | D Rodriguez | 1926 | Bd-GPL | Petrópolis | RJ | -22.50 | -43.18 | Rodriguez et al. 2014 |
| MNRJ 1484 | *Hylodes asper* | Stream | Adult | D Rodriguez | 1928 | Bd-GPL | Petrópolis | RJ | -22.50 | -43.18 | Rodriguez et al. 2014 |

| | | | | | | | | | | | |
|---|---|---|---|---|---|---|---|---|---|---|---|
| MNRJ 1516 | *Hylodes asper* | Stream | Adult | D Rodriguez | 1928 | Bd-GPL | São Jose do Barreiro | SP | -22.64 | -44.56 | Rodriguez et al. 2014 |
| MNRJ 189 | *Cycloramphus semipalmatus* | Stream | Adult | D Rodriguez | 1922 | Bd-GPL | Cubatão | SP | -23.89 | -46.42 | Rodriguez et al. 2014 |
| MNRJ 2142 | *Fritziana fissilis* | Phytotelma | Adult | D Rodriguez | 1951 | Bd-GPL | São Jose do Barreiro | SP | -22.64 | -44.56 | Rodriguez et al. 2014 |
| MNRJ 34713 | *Melanophryniscus moreirae* | Pond | Adult | D Rodriguez | 1949 | Bd-GPL | Petrópolis | RJ | -22.50 | -43.18 | Rodriguez et al. 2014 |
| MNRJ 43610 | *Ololygon ariadne* | Stream | Adult | D Rodriguez | 1951 | Bd-GPL | São Jose do Barreiro | SP | -22.64 | -44.56 | Rodriguez et al. 2014 |
| MNRJ 531 | *Cycloramphus fuliginosus* | Stream | Adult | D Rodriguez | 1923 | Bd-GPL | Campos dos Goytacazes | RJ | -21.74 | -41.33 | Rodriguez et al. 2014 |
| MNRJ 546e | *Hylodes perplicatus* | Stream | Adult | D Rodriguez | 1915 | Bd-GPL | Joinville | SC | -26.30 | -48.84 | Rodriguez et al. 2014 |
| MNRJ 546g | *Hylodes perplicatus* | Stream | Adult | D Rodriguez | 1915 | Bd-GPL | Joinville | SC | -26.30 | -48.84 | Rodriguez et al. 2014 |
| MNRJ 5549 | *Melanophryniscus moreirae* | Pond | Adult | D Rodriguez | 1902 | Bd-GPL | Itatiaia | RJ | -22.35 | -44.63 | Rodriguez et al. 2014 |
| MNRJ 5592 | *Hylodes perplicatus* | Stream | Adult | D Rodriguez | 1916 | Bd-Asia-2/ Brazil | Joinville | SC | -26.30 | -48.84 | Rodriguez et al. 2014 |
| MNRJ 5594 | *Hylodes perplicatus* | Stream | Adult | D Rodriguez | 1916 | Bd-GPL | Joinville | SC | -26.30 | -48.84 | Rodriguez et al. 2014 |
| MNRJ 5607 | *Hylodes perplicatus* | Stream | Adult | D Rodriguez | 1916 | Bd-GPL | Joinville | SC | -26.30 | -48.84 | Rodriguez et al. 2014 |
| MNRJ 5613 | *Hylodes perplicatus* | Stream | Adult | D Rodriguez | 1916 | Bd-GPL | Joinville | SC | -26.30 | -48.84 | Rodriguez et al. 2014 |

| | | | | | | | | | | | |
|---|---|---|---|---|---|---|---|---|---|---|---|
| MNRJ 840 | *Scinax perpusillus* | Phytotelma | Adult | D Rodriguez | 1924 | Bd-GPL | Angra dos Reis | RJ | -23.01 | -44.31 | Rodriguez et al. 2014 |
| MNRJ/ALMN 1390 | *Crossodactylus gaudichaudii* | Stream | Adult | D Rodriguez | 1927 | Bd-GPL | Santo André | SP | -23.78 | -46.30 | Rodriguez et al. 2014 |
| MNRJ/ALMN 1999 | *Crossodactylus gaudichaudii* | Stream | Adult | D Rodriguez | 1929 | Bd-GPL | Rio De janeiro | RJ | -22.90 | -43.20 | Rodriguez et al. 2014 |
| MNRJ/ALMN 2065 | *Crossodactylus dispar* | Stream | Adult | D Rodriguez | 1930 | Bd-GPL | São Jose do Barreiro | SP | -22.64 | -44.56 | Rodriguez et al. 2014 |
| MNRJ/ALMN 2071 | *Crossodactylus dispar* | Stream | Adult | D Rodriguez | 1930 | Bd-GPL | São Jose do Barreiro | SP | -22.64 | -44.56 | Rodriguez et al. 2014 |
| MNRJ/ALMN 480 | *Crossodactylus gaudichaudii* | Stream | Adult | D Rodriguez | 1923 | Bd-Asia-2/ Brazil | Cabo Frio | RJ | -22.86 | -42.04 | Rodriguez et al. 2014 |
| MNRJ/ALMN 87 | *Crossodactylus gaudichaudii* | Stream | Adult | D Rodriguez | 1920 | Bd-GPL | Teresópolis | RJ | -22.46 | -43.00 | Rodriguez et al. 2014 |
| MZUSP 109133 | *Crossodactylus gaudichaudii* | Stream | Adult | D Rodriguez | 1968 | Bd-GPL | São Jose do Barreiro | SP | -22.64 | -44.56 | Rodriguez et al. 2014 |
| MZUSP 109513 | *Crossodactylus bokermanni* | Stream | Adult | D Rodriguez | 1971 | Bd-GPL | Catas Altas | MG | -20.05 | -43.45 | Rodriguez et al. 2014 |
| MZUSP 109645 | *Crossodactylus caramaschii* | Stream | Adult | D Rodriguez | 1958 | Bd-GPL | Florianópolis | SC | -27.49 | -48.40 | Rodriguez et al. 2014 |
| MZUSP 109650 | *Crossodactylus caramaschii* | Stream | Adult | D Rodriguez | 1959 | Bd-GPL | Florianópolis | SC | -27.49 | -48.40 | Rodriguez et al. 2014 |
| MZUSP 109659 | *Crossodactylus caramaschii* | Stream | Adult | D Rodriguez | 1959 | Bd-GPL | Florianópolis | SC | -27.49 | -48.40 | Rodriguez et al. 2014 |
| MZUSP 110332 | *Ololygon ariadne* | Stream | Adult | D Rodriguez | 1968 | Bd-GPL | São Jose do Barreiro | SP | -22.64 | -44.56 | Rodriguez et al. 2014 |

| | | | | | | | | | | | |
|---|---|---|---|---|---|---|---|---|---|---|---|
| MZUSP 110336 | *Ololygon ariadne* | Stream | Adult | D Rodriguez | 1967 | Bd-GPL | São Jose do Barreiro | SP | -22.64 | -44.56 | Rodriguez et al. 2014 |
| MZUSP 110631 | *Crossodactylus bokermanni* | Stream | Adult | D Rodriguez | 1964 | Bd-GPL | Santana do Riacho | MG | -18.96 | -43.71 | Rodriguez et al. 2014 |
| MZUSP 112587 | *Hylodes asper* | Stream | Adult | D Rodriguez | 1965 | Bd-GPL | Marumbi | PR | -23.70 | -51.63 | Rodriguez et al. 2014 |
| MZUSP 112588 | *Hylodes asper* | Stream | Adult | D Rodriguez | 1965 | Bd-GPL | Marumbi | PR | -23.70 | -51.63 | Rodriguez et al. 2014 |
| MZUSP 118 | *Boana pardalis* | Pond | Adult | D Rodriguez | 1942 | Bd-GPL | Itatiaia | RJ | -22.35 | -44.63 | Rodriguez et al. 2014 |
| MZUSP 130376 | *Hylodes perplicatus* | Stream | Adult | D Rodriguez | 2000 | Bd-GPL | Monte Alegre dos Campos | RS | -28.78 | -50.79 | Rodriguez et al. 2014 |
| MZUSP 136242 | *Fritziana fissilis* | Phytotelma | Adult | D Rodriguez | 2006 | Hybrid or Coinfection | Bertioga | SP | -23.71 | -46.03 | Rodriguez et al. 2014 |
| MZUSP 209 | *Bokermannohyla luctuosa* | Pond | Adult | D Rodriguez | 1897 | Bd-GPL | Piquete | SP | -22.61 | -45.17 | Rodriguez et al. 2014 |
| MZUSP 240 | *Boana faber* | Pond | Adult | D Rodriguez | 1907 | Bd-GPL | Santo André | SP | -23.78 | -46.30 | Rodriguez et al. 2014 |
| MZUSP 296 | *Scinax hayii* | Pond | Adult | D Rodriguez | 1902 | Bd-GPL | Campos do Jordão | SP | -22.73 | -45.59 | Rodriguez et al. 2014 |
| MZUSP 53055 | *Vitreorana eurygnatha* | Stream | Adult | D Rodriguez | 1976 | Bd-GPL | Itapeva | SP | -23.96 | -48.90 | Rodriguez et al. 2014 |
| MZUSP 53058 | *Vitreorana eurygnatha* | Stream | Adult | D Rodriguez | 1976 | Bd-GPL | Itapeva | SP | -23.96 | -48.90 | Rodriguez et al. 2014 |
| MZUSP 53330 | *Megaelosia goeldii* | Stream | Adult | D Rodriguez | 1977 | Bd-GPL | Teresópolis | RJ | -22.46 | -43.00 | Rodriguez et al. 2014 |

| | | | | | | | | | | | |
|---|---|---|---|---|---|---|---|---|---|---|---|
| MZUSP 56836 | *Crossodactylus bokermanni* | Stream | Adult | D Rodriguez | 1979 | Bd-Asia-2/ Brazil | Santana do Riacho | MG | -18.96 | -43.71 | Rodriguez et al. 2014 |
| MZUSP 58677 | *Hylodes heyeri* | Stream | Adult | D Rodriguez | 1972 | Bd-GPL | Iporanga | SP | -24.48 | -48.65 | Rodriguez et al. 2014 |
| MZUSP 58744 | *Cycloramphus boraceiensis* | Stream | Adult | D Rodriguez | 1982 | Bd-Asia-2/ Brazil | São Sebastiao | SP | -23.76 | -45.41 | Rodriguez et al. 2014 |
| MZUSP 60698 | *Hylodes perplicatus* | Stream | Adult | D Rodriguez | 1982 | Bd-GPL | São Sebastiao | SP | -23.76 | -45.41 | Rodriguez et al. 2014 |
| MZUSP 64712 | *Phyllomedusa distincta* | Pond | Adult | D Rodriguez | 1949 | Bd-GPL | São Bento do Sul | SC | -26.24 | -49.38 | Rodriguez et al. 2014 |
| MZUSP 64714 | *Phyllomedusa distincta* | Pond | Adult | D Rodriguez | 1949 | Bd-GPL | São Bento do Sul | SC | -26.24 | -49.38 | Rodriguez et al. 2014 |
| MZUSP 75694 | *Aplastodiscus perviridis* | Stream/ Pond | Adult | D Rodriguez | 1970 | Bd-GPL | Botucatu | SP | -22.89 | -48.45 | Rodriguez et al. 2014 |
| MZUSP 76436 | *Fritziana goeldii* | Phytotelma | Adult | D Rodriguez | 1964 | Bd-GPL | Teresópolis | RJ | -22.46 | -43.00 | Rodriguez et al. 2014 |
| MZUSP 76695 | *Proceratophrys boiei* | Stream/ Pond | Adult | D Rodriguez | 1964 | Bd-GPL | Teresópolis | RJ | -22.46 | -43.00 | Rodriguez et al. 2014 |
| MZUSP 77615 | *Crossodactylus bokermanni* | Stream | Adult | D Rodriguez | 1973 | Bd-GPL | Santana do Riacho | MG | -18.96 | -43.71 | Rodriguez et al. 2014 |
| MZUSP 86 | *Boana prasina* | Stream/ Pond | Adult | D Rodriguez | 1905 | Bd-GPL | Campos do Jordão | SP | -22.73 | -45.59 | Rodriguez et al. 2014 |

* CLFT: Coleção de Culturas Luís Felipe Toledo; MZUSP: Museu de Zoologia, Universidade de São Paulo; MNRJ: Museu Nacional, Rio de Janeiro; AL: state of Alagoas; BA: state of Bahia; ES: state of Espírito Santo; MG: state of Minas Gerais; PR: state of Paraná; RJ: state of Rio de Janeiro; SC: state of Santa Catarina; SP: state of São Paulo.

**Table S2.** *Batrachochytrium dendrobatidis* (Bd) genotypes used in the challenge assay. The table includes isolate/genotype name, designation (panzootic 'P', enzootic 'E' and hybrid 'H'), chytrid lineages or genotype, locality (municipality, state), host species, year of isolation, and number of passages for each isolate.

| Isolate | Designation | Lineages or genotype | Locality | Host species | Year | Passage |
|---|---|---|---|---|---|---|
| CLFT 168 | P1 | Bd-GPL | Alto Caparaó, MG | *Phantasmarana apuana* | 2015 | 7 |
| CLFT 198 | P2 | Bd-GPL | Santa Isabel, SP | *Aquarana catesbeiana* | 2016 | 4 |
| CLFT 041 | E1 | Bd-Asia-2/ Brazil | Morretes, PR | *Bokermannohyla hylax* | 2013 | 15 |
| CLFT 172 | E2 | Bd-Asia-2/ Brazil | Pindamonhangaba , SP | *Aquarana catesbeiana* | 2016 | 5 |
| CLFT 024-02 | H | Hybrid | Morretes, PR | *Hylodes cardosoi* | 2011 | 20 |

**Table S3.** Treatment groups of the infection assay with five different genotypes of the fungus *Batrachochytrium dendrobatidis* (Bd), which included single-genotype and mixed genotype exposures with either two or three genotypes in all possible combinations, and a control group. Designations (panzootic 'P', enzootic 'E' and hybrid 'H'), number of hosts per treatment, and chytrid lineages or genotype.

| Treatment Groups | n | Lineages or genotype |
|---|---|---|
| P1 | 16 | Bd-GPL |
| P2 | 16 | Bd-GPL |
| E1 | 16 | Bd-Asia-2/Brazil |
| E2 | 16 | Bd-Asia-2/Brazil |
| H | 16 | Hybrid |
| P1 + E1 | 15 | Bd-GPL + Bd-Asia-2/Brazil |
| P1 + E2 | 15 | Bd-GPL + Bd-Asia-2/Brazil |
| P1 + H | 14 | Bd-GPL + Hybrid |
| P2 + E1 | 14 | Bd-GPL + Bd-Asia-2/Brazil |
| P2 + E2 | 16 | Bd-GPL + Bd-Asia-2/Brazil |
| P2 + H | 15 | Bd-GPL + Hybrid |
| E2 + H | 16 | Bd-Asia-2/Brazil + Hybrid |
| E1 + H | 16 | Bd-Asia-2/Brazil + Hybrid |
| P1 + E1 + H | 16 | Bd-GPL + Bd-Asia-2/Brazil + Hybrid |
| P1 + E2 + H | 16 | Bd-GPL + Bd-Asia-2/Brazil + Hybrid |
| P2 + E1 + H | 16 | Bd-GPL + Bd-Asia-2/Brazil + Hybrid |
| P2 + E2 + H | 16 | Bd-GPL + Bd-Asia-2/Brazil + Hybrid |
| Control | 32 | Control |

**Table S4.** Applied Biosystems assay information for BdSC9_621917_AC (Assay ID AHBKG6X) at 40X concentration.

| Primer/Probe | Sequence | Concentration (μM) | Reporter | Quencher |
|---|---|---|---|---|
| Forward Primer | GAGCTGGCCTTTCTCTTGAGA | 36 | | |
| Reverse Primer | CGTAGAATAAGAAGACATTGCACTTGGT | 36 | | |
| Reporter 1 | AGATCAAAATGGTCACTCAT | 8 | VIC | NFQ |
| Reporter 2 | CAAAATGGGCACTCAT | 8 | FAM | NFQ |

**Table S5**. Standard curve parameters for assay Bdmt_26360_AC, when 1.34E-5 ng/µl, 1.34E-4 ng/µl, 0.00134 ng/µl, 0.0134 ng/µl, 0.134 ng/µl of DNA were used as input for strains BAF01 (Bd-Asia-2/Brazil), NAF01 (*Bd*-GPL), and CLFT024-02 (hybrid). Reactions were run in triplicate.

| Reporter | Allele | Target Genome | Slope | Y-Intercept | $R^2$ | Efficiency (%) | Std Dev | Std Error |
|---|---|---|---|---|---|---|---|---|
| FAM | A | BAF 01 | -3.4276 | 20.5516 | 0.9984 | 95.7714 | 0.0330 | 0.0506 |
| VIC | G | NAF 01 | -3.2988 | 21.4904 | 0.9930 | 100.9743 | 0.1757 | 0.0765 |
| VIC | G | CLFT 024-02 | -3.4318 | 20.2440 | 0.9992 | 95.6105 | 0.0192 | 0.0377 |

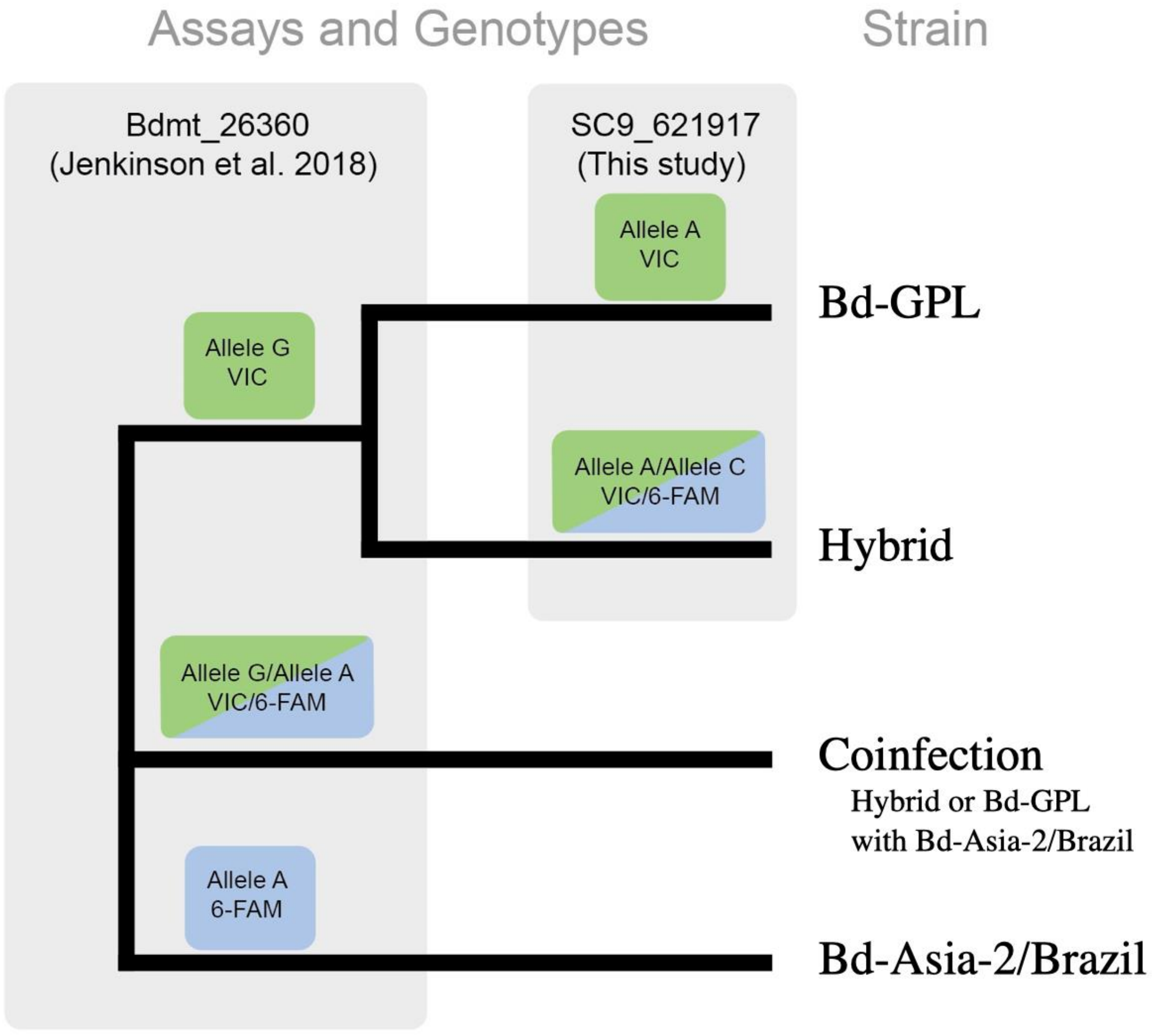


**Figure S1.** SNPASSAY. Diagram of diagnostic genotyping assays used in this study.

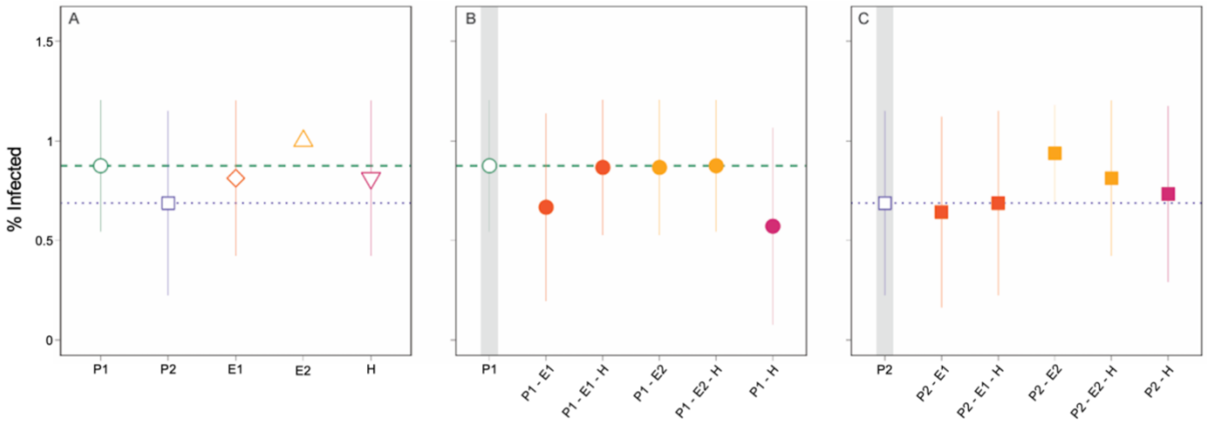


**Figure S2.** The presence of multiple genotypes did not affect the proportion of hosts infected (Binomial GLM, all *p*-values > 0.05). Grey bars highlight single-genotype infections. These results serve as a quality control metric and clarify that our infection methodology was consistent across treatments.

**Dataset S1.** Sequence data in fasta format used to explore the phylogenetic relationships between the *Bd* isolates used in this study.

**>CLFT024**
CGCATGGCAGAGGCAAGATGGCAGCGCGAATGTCARCAGCTTGTTCAAAAATTAGA
ATACGAGGTAGATGGTTTAAGCCTGCTAAATTTACCACAACAAAAATCTAATATGCA
TGATATAGTTGAACAGGTCAGATGAGTTGCAGCGTCTTTTCGATCAGCAAGTGCTCA
AGACACGGGAATTAGAGAGRGAGCTGGCTGACACCCGACTGCTTTTGCACCAGACT
CAATCTGGTTAGTGCAGTGTTTACTTTTTAACTATTTGTCGTTGAATTTCATTCGAAT
TAACTTATTTATCTGATGCGTGGAGGGTATTACCACGAAGTTCGGCTTCACGGATAG
TACGAGCCAAATTGTCACCTGGAAGAGCCTAGTAGAGATCATTGATTACAACATTCA
ACAATGAAGGGTTAGTATGCGCGATTTACTACGACTGAGCTGATGCATCATGTAATA
GGATCAGCAAAAGTATCTGCAGATGCAATAAACCCATTAACAATGCTTAAAGATCA
TTAAGCAGTGATATCCGTTATATTTCAACAACTGCTGATCAGTCATACAAGTCAATG
CAATCAACGAATGRCAGCATCAGTTGTACTAACAACTGATGCTGAATTGATCAAGA
GTAAAGACTACCGCAAAGCATTGTTTTAAACAAGACATGCTCTCCACTATCCTCATG
ATAATCAAGCCGTCAAAGACACCAATAGCAATGAGTCTGTGCGCAGGATAAAAAAC
CCTTCACCCACTGTGTATCAATGCATTCTTACTCTAAGAAGACAAGCCTTGAGCATA
TGAGCAGTTCCAGGAGCATCAGCAGTGTCGAAGCGAGAGCCAGCATTGACCACAAT
TGCCAGAGTCGAAGCAGGCCCAAAGTGGTCGTACGTGGCCACATTGACATGATGGT
TCGTTTCCATGGATYATCAGTATGTRTATCTGCTGGATTGGACTGTATCGGGGAATG
TATGGTTCTATGTGTATTAGACATTGGAGAAACTGGAGTTGACGATCGTTGTGACGG
AGTTGAAGGTTTTTTACGAGATGCATTCGTAGTCTGTAAGCATCGAGCAACCCAAGG
CATAGTAGAATCAATCTGTATTTCAGCAGTCCATCCACCCATTGATCGATCTGTCTTC
CATTGCAAAGACGGTTTACCAAAATGTGATAGCAGCTGATCCAAGAGAATCATTCCT
ACAGAGTGTCGGGTTGTTGGATAGGTAAAGTTTCCCAATCCAACCACCTGTCATTGC
AGCGATTGAATCAATAGTTAGCTGGATGGTTCACACTGATGACTTGAATATCAACTA
TTCTGACTTGCCAATACAGGAGATAGTTGTTTATACTGCAACTCGGTAGTTACAGAT
GGAATAGTAGAATAACTAGCAGTGGAGAATGGCTGGCCTCGCATTGATACAAAACC
ATATACCAAGACCGAATTATGGTTAGTATATATTATTCGCTGCATTGAGTTTTTAGTT
GGCTTGTACCAATCTAACAAKGCTGTGTATGAAGCTATCATTCTGAAAATCGGAAAT
TTGATTCAATCAAGGCTCACGTTGGTTTGCAAATTGACGAAATTCAACTCAGGRTTA
TGCTAAATGATAGCGAGGTTAGTATTGTTRTGTTGTTGCATTTACAATGTTTGTTATA
CACTAATGTCTTGRATATTATGYATATCAGGTTATTGGTGTCAAGGATAGCAGTAAA
TGGAATTGGGATATTATTGAAGAAATAGTATCTGGTCCACTTTTGAATTCAAAACGC
ATTCACTACTCATTTCAAGCCACCTGTGYTGGGTTTATTTGCACGAGTAATGGGTATT
CGTAAAGCCAGTGGACGGTCTCCGTCAACCGCTTTGACCCAGGATATTTCTCCGGGT
AATTGTTGGGCCATGTCAGGTAAAATTAAATTCAAGTGGTGAAAATGTGTTGATGTT
TGGATAAACAATGTTTGATAATTTGTTCTACCCGTGACATGATGATTAGACTCGAGC
GGCACGTTGGCGATATCCCTTGCCGAGCCCATCATACCAACTGATATGACCATTGAG
CATTCTCATATCCAAACGTCCATTRGTGACCGACACACATCAGCTCCACGCCAAATC
GAACTTTGGGCAGTCTTTGATGCTGTAGAATTTGCTAAACTTGATCTAAACAACAAT
CAGGTCAGACTAC

**>CLFT041**

CGCATGGCAGAGGCAAGATGGCAGCGCGAATGTCAACAGCTTGTTCAAAAATTAGA
ATACGAGGTAGATGGTTTAAGCCTGCTAAATTTACCACAACAAAAATCTAATATGCA
TGATATAGTTGAACAGGTCAGATGAGTTGCAGCGTCTTTTCGATCAGCAAGTGCTCA
AGACACGGGAATTAGAGAGGGAGCTGGCTGACACCCGACTGCTTTTGCACCAGACT
CAATCTGGTTAGTGCAGTGTTTACTTTTTAACTATTTGTCGTTGAATTTCATTCGAAT
TAACTTATTTATCTGATGCGTGGAGGGTATTACCACGAAGTTCGGCTTCACGGATAG
TACGAGCCAAATTGTCACCTGGAAGAGCCTAGTAGAGATCATTGATTACAACATTCA
ACAATGAAGGGTTAGTATGCGCGATTTACTACGACTGAGCTGATGCATCATGTAATA
GGATCAGCAAAAGTATCTGCAGATGCAATAAACCCATTAACAATGCTTAAAGATCA
TTAAGCAGTGATATCCGTTATATTTCAACAACTGCTGATCAGTCATACAAGTCAATG
CAATCAACGAATGGCAGCATCAGTTGTACTAACAACTGATGCTGAATTGATCAAGA
GTAAAGACTACCGCAAAGCATTGTTTTAAACAAGACATGCTCTCCACTATCCTCATG
ATAATCAAGCCGTCAAAGACACCAATAGCAATGAGTCTGTGCGCAGGATAAAAAAC
CCTTCACCCACTGTGTATCAATGCATTCTTACTCTAAGAAGACAAGCCTTGAGCATA
TGAGCAGTTCCAGGAGCATCAGCAGTGTCGAAGCGAGAGCCAGCATTGACCACAAT
TGCCAGAGTCGAAGCAGGCCCAAAGTGGTCGTACGTGGCCACATTGACATGATGGT
TCGTTTCCATGGATTATCAGTATGTATATCTGCTGGATTGGACTGTATCGGGGAATGT
ATGGTTCTATGTGTATTAGACATTGGAGAAACTGGAGTTGACGATCGTTGTGACGGA
GTTGAAGGTTTTTTACGAGATGCATTCGTAGTCTGTAAGCATCGAGCAACCCAAGGC
ATAGTAGAATCAATCTGTATTTCAGCAGTCCATCCACCCATTGATCGATCTGTCTTCC
ATTGCAAAGACGGTTTACCAAAATGTGATAGCAGCTGATCCAAGAGAATCATTCCTA
CAGAGTGTCGGGTTGTTGGATAGGTAAAGTTTCCCAATCCAACCACCTGTCATTGCA
GCGATTGAATCAATAGTTAGCTGGATGGTTCACACTGATGACTTGAATATCAACTAT
TCTGACTTGCCAATACAGGAGATAGTTGTTTATACTGCAACTCGGTAGTTACAGATG
GAATAGTAGAATAACTAGCAGTGGAGAATGGCTGGCCTCGCATTGATACAAAACCA
TATACCAAGACCGAATTATGGTTAGTATATATTATTCGCTGCATTGAGTTTTTAGTTG
GCTTGTAYCAATCTAACAAKGCTGTGTATGAAGCTATCATTCTGAAAATCGGAAATT
TGATTCAATCAAGGCTCACGTTGGTTTGCAAATTGACGAAATTCAACTCAGGGTTAT
GCTAAATGATAGCGAGGTTAGTATTGTTRTGTTGTTGCATTTACAATGTTTGTTATAC
ACTAATGTCTTGGATATTATGCATATCAGGTTATTGGTGTCAAGGATAGCAGTAAAT
GGAATTGGGATATTATTGAAGAAATAGTATCTGGTCCACTTTTGAATTCAAAACGCA
TTCACTACTCATTTCAAGCCACCTGTGTTGGGTTTATTTGCACGAGTAATGGGTATTC
GTAAAGCCAGTGGACGGTCTCCGTCAACCGCTTTGACCCAGGATATTTCTCCGGGTA
ATTGTTGGGCCATGTCAGGTAAAATTAAATTCAAGTGGTGAAAATGTGTTGATGTTT
GGATAAACAATGTTTGATAATTTGTTCTACCCGTGACATGATGATTAGACTCGAGCG
GCACGTTGGCGATATCCCTTGCCGAGCCCATCATACCAACTGATATGACCATTGAGC
ATTCTCATATCCAAACGTCCATTGGTGACCGACACACATCAGCTCCACGCCAAATCG
AACTTTGGGCAGTCTTTGATGCTGTAGAATTTGCTAAACTTGATCTAAACAACAATC
AGGTCAGACTAC

**>CLFT172**
CGCATGGCAGAGGCAAGATGGCAGCGCGAATGTCAACAGCTTGTTCAAAAATTAGA
ATACGAGGTAGATGGTTTAAGCCTGCTAAATTTACCACAACAAAAATCTAATATGCA
TGATATAGTTGAACAGGTCAGATGAGTTGCAGCGTCTTTTCGATCAGCAAGTGCTCA
AGACACGGGAATTAGAGAGGGAGCTGGCTGACACCCGACTGCTTTTGCACCAGACT
CAATCTGGTTAGTGCAGTGTTTACTTTTTAACTATTTGTCGTTGAATTTCATTCGAAT

TAACTTATTTATCTGATGCGTGGAGGGTATTACCACGAAGTTCGGCTTCACGGATAG
TACGAGCCAAATTGTCACCTGGAAGAGCCTAGTAGAGATCATTGATTACAACATTCA
ACAATGAAGGGTTAGTATGCGCGATTTACTACGACTGAGCTGATGCATCATGTAATA
GGATCAGCAAAAGTATCTGCAGATGCAATAAACCCATTAACAATGCTTAAAGATCA
TTAAGCAGTGATATCCGTTATATTTCAACAACTGCTGATCAGTCATACAAGTCAATG
CAATCAACGAATGGCAGCATCAGTTGTACTAACAACTGATGCTGAATTGATCAAGA
GTAAAGACTACCGCAAAGCATTGTTTTAAACAAGACATGCTCTCCACTATCCTCATG
ATAATCAAGCCGTCAAAGACACCAATAGCAATGAGTCTGTGCGCAGGATAAAAAAC
CCTTCACCCACTGTGTATCAATGCATTCTTACTCTAAGAAGACAAGCCTTGAGCATA
TGAGCAGTTCCAGGAGCATCAGCAGTGTCGAAGCGAGAGCCAGCATTGACCACAAT
TGCCAGAGTCGAAGCAGGCCCAAAGTGGTCGTACGTGGCCACATTGACATGATGGT
TCGTTTCCATGGATTATCAGTATGTATATCTGCTGGATTGGACTGTATCGGGGAATGT
ATGGTTCTATGTGTATTAGACATTGGAGAAACTGGAGTTGACGATCGTTGTGACGGA
GTTGAAGGTTTTTTACGAGATGCATTCGTAGTCTGTAAGCATCGAGCAACCCAAGGC
ATAGTAGAATCAATCTGTATTTCAGCAGTCCATCCACCCATTGATCGATCTGTCTTCC
ATTGCAAAGACGGTTTACCAAAATGTGATAGCAGCTGATCCAAGAGAATCATTCCTA
CAGAGTGTCGGGTTGTTGGATAGGTAAAGTTTCCCAATCCAACCACCTGTCATTGCA
GCGATTGAATCAATAGTTAGCTGGATGGTTCACACTGATGACTTGAATATCAACTAT
TCTGACTTGCCAATACAGGAGATAGTTGTTTATACTGCAACTCGGTAGTTACAGATG
GAATAGTAGAATAACTAGCAGTGGAGAATGGCTGGCCTCGCATTGATACAAAACCA
TATACCAAGACCGAATTATGGTTAGTATATATTATTCGCTGCATTGAGTTTTTAGTTG
GCTTGTATCAATCTAACAATGCTGTGTATGAAGCTATCATTCTGAAAATCGGAAATT
TGATTCAATCAAGGCTCACGTTGGTTTGCAAATTGACGAAATTCAACTCAGGGTTAT
GCTAAATGATAGCGAGGTTAGTATTGTTATGTTGTTGCATTTACAATGTTTGTTATAC
ACTAATGTCTTGGATATTATGCATATCAGGTTATTGGTGTCAAGGATAGCAGTAAAT
GGAATTGGGATATTATTGAAGAAATAGTATCTGGTCCACTTTTGAATTCAAAACGCA
TTCACTACTCATTTCAAGCCACCTGTGTTGGGTTTATTTGCACGAGTAATGGGTATTC
GTAAAGCCAGTGGACGGTCTCCGTCAACCGCTTTGACCCAGGATATTTCTCCGGGTA
ATTGTTGGGCCATGTCAGGTAAAATTAAATTCAAGTGGTGAAAATGTGTTGATGTTT
GGATAAACAATGTTTGATAATTTGTTCTACCCGTGACATGATGATTAGACTCGAGCG
GCACGTTGGCGATATCCCTTGCCGAGCCCATCATACCAACTGATATGACCATTGAGC
ATTCTCATATCCAAACGTCCATTGGTGACCGACACACATCAGCTCCACGCCAAATCG
AACTTTGGGCAGTCTTTGATGCTGTAGAATTTGCTAAACTTGATCTAAACAACAATC
AGGTCAGACTAC

**>CLFT168**
CGCATGGCAGAGGCAAGATGGCAGCGCGAATGTCAGCAGCTTGTTCAAAAATTAGA
ATACGAGGTAGATGGTTTAAGCCTGCTAAATTTACCACAACAAAAATCTAATATGCA
TGATATAGTTGAACAGGTCAGATGAGTTGCAGCGTCTTTTCGATCAGCAAGTGCTCA
AGACACGGGAATTAGAGAGAGAGCTGGCTGACACCCGACTGCTTTTGCACCAGACT
CAATCTGGTTAGTGCAGTGTTTACTTTTTAACTATTTGTCGTTGAATTTCATTCGAAT
TAACTTATTTATCTGATGCGTGGAGGGTATTACCACGAAGTTCGGCTTCACGGATAG
TACGAGCCAAATTGTCACCTGGAAGAGCCTAGTAGAGATCATTGATTACAACATTCA
ACAATGAAGGGTTAGTATGCGCGATTTACTACGACTGAGCTGATGCATCATGTAATA
GGATCAGCAAAAGTATCTGCAGATGCAATAAACCCATTAACAATGCTTAAAGATCA
TTAAGCAGTGATATCCGTTATATTTCAACAACTGCTGATCAGTCATACAAGTCAATG

CAATCAACGAATGACAGCATCAGTTGTACTAACAACTGATGCTGAATTGATCAAGA
GTAAAGACTACCGCAAAGCATTGTTTTAAACAAGACATGCTCTCCACTATCCTCATG
ATAATCAAGCCGTCAAAGACACCAATAGCAATGAGTCTGTGCGCAGGATAAAAAAC
CCTTCACCCACTGTGTATCAATGCATTCTTACTCTAAGAAGACAAGCCTTGAGCATA
TGAGCAGTTCCAGGAGCATCAGCAGTGTCGAAGCGAGAGCCAGCATTGACCACAAT
TGCCAGAGTCGAAGCAGGCCCAAAGTGGTCGTACGTGGCCACATTGACATGATGGT
TCGTTTCCATGGATCATCAGTATGTGTATCTGCTGGATTGGACTGTATCGGGGAATG
TATGGTTCTATGTGTATTAGACATTGGAGAAACTGGAGTTGACGATCGTTGTGACGG
AGTTGAAGGTTTTTTACGAGATGCATTCGTAGTCTGTAAGCATCGAGCAACCCAAGG
CATAGTAGAATCAATCTGTATTTCAGCAGTCCATCCACCCATTGATCGATCTGTCTTC
CATTGCAAAGACGGTTTACCAAAATGTGATAGCAGCTGATCCAAGAGAATCATTCCT
ACAGAGTGTCGGGTTGTTGGATAGGTAAAGTTTCCCAATCCAACCACCTGTCATTGC
AGCGATTGAATCAATAGTTAGCTGGATGGTTCACACTGATGACTTGAATATCAACTA
TTCTGACTTGCCAATACAGGAGATAGTTGTTTATACTGCAACTCGGTAGTTACAGAT
GGAATAGTAGAATAACTAGCAGTGGAGAATGGCTGGCCTCGCATTGATACAAAACC
ATATACCAAGACCGAATTATGGTTAGTATATATTATTCGCTGCATTGAGTTTTTAGTT
GGCTTGTACCAATCTAACAATGCTGTGTATGAAGCTATCATTCTGAAAATCGGAAAT
TTGATTCAATCAAGGCTCACGTTGGTTTGCAAATTGACGAAATTCAACTCAGGATTA
TGCTAAATGATAGCGAGGTTAGTATTGTTATGTTGTTGCATTTACAATGTTTGTTATA
CACTAATGTCTTGAATATTATGTATATCAGGTTATTGGTGTCAAGGATAGCAGTAAA
TGGAATTGGGATATTATTGAAGAAATAGTATCTGGTCCACTTTTGAATTCAAAACGC
ATTCACTACTCATTTCAAGCCACCTGTGCTGGGTTTATTTGCACGAGTAATGGGTATT
CGTAAAGCCAGTGGACGGTCTCCGTCAACCGCTTTGACCCAGGATATTTCTCCGGGT
AATTGTTGGGCCATGTCAGGTAAAATTAAATTCAAGTGGTGAAAATGTGTTGATGTT
TGGATAAACAATGTTTGATAATTTGTTCTACCCGTGACATGATGATTAGACTCGAGC
GGCACGTTGGCGATATCCCTTGCCGAGCCCATCATACCAACTGATATGACCATTGAG
CATTCTCATATCCAAACGTCCATTAGTGACCGACACACATCAGCTCCACGCCAAATC
GAACTTTGGGCAGTCTTTGATGCTGTAGAATTTGCTAAACTTGATCTAAACAACAAT
CAGGTCAGACTAC

**>CLFT198**
CGCATGGCAGAGGCAAGATGGCAGCGCGAATGTCAGCAGCTTGTTCAAAAATTAGA
ATACGAGGTAGATGGTTTAAGCCTGCTAAATTTACCACAACAAAAATCTAATATGCA
TGATATAGTTGAACAGGTCAGATGAGTTGCAGCGTCTTTTCGATCAGCAAGTGCTCA
AGACACGGGAATTAGAGAGAGAGCTGGCTGACACCCGACTGCTTTTGCACCAGACT
CAATCTGGTTAGTGCAGTGTTTACTTTTTAACTATTTGTCGTTGAATTTCATTCGAAT
TAACTTATTTATCTGATGCGTGGAGGGTATTACCACGAAGTTCGGCTTCACGGATAG
TACGAGCCAAATTGTCACCTGGAAGAGCCTAGTAGAGATCATTGATTACAACATTCA
ACAATGAAGGGTTAGTATGCGCGATTTACTACGACTGAGCTGATGCATCATGTAATA
GGATCAGCAAAAGTATCTGCAGATGCAATAAACCCATTAACAATGCTTAAAGATCA
TTAAGCAGTGATATCCGTTATATTTCAACAACTGCTGATCAGTCATACAAGTCAATG
CAATCAACGAATGACAGCATCAGTTGTACTAACAACTGATGCTGAATTGATCAAGA
GTAAAGACTACCGCAAAGCATTGTTTTAAACAAGACATGCTCTCCACTATCCTCATG
ATAATCAAGCCGTCAAAGACACCAATAGCAATGAGTCTGTGCGCAGGATAAAAAAC
CCTTCACCCACTGTGTATCAATGCATTCTTACTCTAAGAAGACAAGCCTTGAGCATA
TGAGCAGTTCCAGGAGCATCAGCAGTGTCGAAGCGAGAGCCAGCATTGACCACAAT

TGCCAGAGTCGAAGCAGGCCCAAAGTGGTCGTACGTGGCCACATTGACATGATGGT
TCGTTTCCATGGATCATCAGTATGTRTATCTGCTGGATTGGACTGTATCGGRGAATGT
ATGGTTCTATGTGYATTAGACATTGGAGAAACTGGAGTTGACGATCGTTGTGACGGA
GTTGAAGGTTTTTTACGAGATGCATTCGTAGTCTGTAAGCATCGAGCAACCCAAGGC
ATAGTAGAATCAATCTGTATTTCAGCAGTCCATCCACCCATTGATCGATCTGTCTTCC
ATTGCAAAGACGGTTTACCAAAATGTGATAGCAGCTGATCCAAGAGAATCATTCCTA
CAGAGTGTCGGGTTGTTGGATAGGTAAAGTTTCCCAATCCAACCACCTGTCATTGCA
GCGATTGAATCAATAGTTAGCTGGATGGTTCACACTGATGACTTGAATATCAACTAT
TCTGACTTGCCAATACAGGAGATAGTTGTTTATACTGCAACTCGGTAGTTACAGATG
GAATAGTAGAATAACTAGCAGTGGAGAATGGCTGGCCTCGCATTGATACAAAACCA
TATACCAAGACCGAATTATGGTTAGTATATATTATTCGCTGCATTGAGTTTTTAGTTG
GCTTGTACCAATCTAACAATGCTGTGTATGAAGCTATCATTCTGAAAATCGGAAATT
TGATTCAATCAAGGCTCACGTTGGTTTGCAAATTGACGAAATTCAACTCAGGATTAT
GCTAAATGATAGCGAGGTTAGTATTGTTATGTTGTTGCATTTACAATGTTTGTTATAC
ACTAATGTCTTGAATATTATGTATATCAGGTTATTGGTGTCAAGGATAGCAGTAAAT
GGAATTGGGATATTATTGAAGAAATAGTATCTGGTCCACTTTTGAATTCAAAACGCA
TTCACTACTCATTTCAAGCCACCTGTGCTGGGTTTATTTGCACGAGTAATGGGTATTC
GTAAAGCCAGTGGACGGTCTCCGTCAACCGCTTTGACCCAGGATATTTCTCCGGGTA
ATTGTTGGGCCATGTCAGGTAAAATTAAATTCAAGTGGTGAAAATGTGTTGATGTTT
GGATAAACAATGTTTGATAATTTGTTCTACCCGTGACATGATGATTAGACTCGAGCG
GCACGTTGGCGATATCCCTTGCCGAGCCCATCATACCAACTGATATGACCATTGAGC
ATTCTCATATCCAAACGTCCATTAGTGACCGACACACATCAGCTCCACGCCAAATCG
AACTTTGGGCAGTCTTTGATGCTGTAGAATTTGCTAAACTTGATCTAAACAACAATC
AGGTCAGACTAC
